# Insulin-like Growth Factor-1 Synergizes with IL-2 to Induce Homeostatic Proliferation of Regulatory T cells

**DOI:** 10.1101/2022.05.12.491665

**Authors:** Melanie R. Shapiro, Leeana D. Peters, Matthew E. Brown, Cecilia Cabello-Kindelan, Amanda L. Posgai, Allison L. Bayer, Todd M. Brusko

## Abstract

IL-2 has been proposed to restore tolerance via regulatory T cell (Treg) expansion in autoimmunity, yet off-target effects necessitate identification of a combinatorial approach. We recently reported reduced levels of immunoregulatory insulin-like growth factor-1 (IGF1) during type 1 diabetes (T1D) progression. Thus, we hypothesized that IGF1 would synergize with IL-2 to expand Tregs. We observed IGF1R was elevated on murine memory and human naïve Treg subsets. IL-2 and IGF1 promoted murine PI3K/Akt and human STAT5 signaling in Tregs. IL-2 and IGF1 treatment expanded Tregs beyond either agent alone in NOD mice. Incubation of naïve human CD4^+^ T cells with IL-2 and IGF1 enhanced Treg proliferation *in vitro*, without the need for T cell receptor ligation. This synergism was attributed to increased high-affinity IL-2Rα expression on naïve Tregs, in contrast to intermediate-affinity IL-2Rβ and IL-2Rγ subunit enhancement on naïve conventional T cells (Tconv). We then demonstrated that IGF1 and IL-2 or the IL2Rγ-chain-dependent cytokine, IL-7, can be used to induce proliferation of genetically-engineered naïve Treg or Tconv cells, respectively. These data support the potential use of IGF1 in combination with common γ-chain cytokines to drive T cell expansions both *in vitro* and *in vivo* for cellular therapeutics and genetic modifications.

## INTRODUCTION

Impairments in immunoregulation— including defective regulatory T cell (Treg) generation as well as peripheral maintenance of phenotypically stable and functional Treg populations— have been suggested to contribute to the development of type 1 diabetes (T1D) and other autoimmune diseases (1, 2). While human studies have largely shown that Treg numbers are not deficient in the peripheral blood of subjects with T1D or other organ-specific autoimmune conditions (1, 3–5), a dearth in the frequency of Tregs at the site of autoimmunity appears to be unique to T1D (6, 7). In contrast, defects in Treg suppressive function have been widely reported in multiple autoimmune conditions (1, 4), including T1D, wherein maintenance of Treg function is compromised in pancreatic draining lymph nodes (pLN) of affected human organ donors (6). Efforts to specifically increase Treg without influencing effector T cell (Teff) numbers are therefore of great interest for T1D prevention.

Low-dose IL-2 has been proposed as a means to selectively enhance the proliferation and function of Tregs, which constitutively express CD25 (IL-2Rα), the alpha chain subunit that creates the high-affinity trimeric IL-2 receptor (IL-2R) complex. This is in direct contrast to conventional CD4^+^ T cells (Tconv), which must upregulate CD25 following activation (8). IL-2Rα pairs with the intermediate-affinity subunits IL-2Rβ and IL-2Rγ, which are less prone to change their expression with activation (9). While the IL-2Rγ subunit, also known as the common-γ chain, can pair with several other cytokine receptor subunits to facilitate signaling by a wide variety of cytokines including IL-7, 9, 15, and 21 in both Treg and Tconv (10), the restricted use of IL-2Rα in IL-2R belies the Treg-specific dependence on IL-2.

Impaired IL-2 signaling has been reported in Tregs of T1D subjects (11, 12). Moreover, decreased STAT5 phosphorylation downstream of IL-2R has been associated with T1D-risk single nucleotide polymorphisms (SNPs) in *CD25* and *PTPN2*, a phosphatase which inhibits JAK1 and JAK3 in the IL-2R pathway (12–14). Similarly, the NOD mouse carries a risk locus containing the *Il2* gene (15) thought to contribute to impaired Stat5 phosphorylation (16) and diminished survival of intra-islet Tregs (17). These defects have been corrected with low-dose IL-2 therapy in both mice (17) and humans (18), with success in improving IL-2 targeting to murine Tregs via the addition of an IL-2-specific monoclonal antibody (IAC: IL-2 Antibody Complex) enhancing IL-2 half-life and blocking interaction with the intermediate-affinity IL-2R complex (19). Despite such improvements to IL-2 delivery, inflammatory cell subsets expressing CD25, such as memory T cells and natural killer (NK) cells, have been reported to expand alongside Tregs (18–20). This highlights the need to identify combination therapies that synergize with IL-2, allowing for efficacy with lower IL-2 doses and avoidance of off-target effects.

Insulin-like growth factor-1 (IGF1) is an immunoregulatory hormone (21) that we recently showed to be significantly reduced in serum from individuals during pre-T1D and following T1D diagnosis (22), potentially contributing to the progression of autoimmunity (23). Low peripheral IGF1 levels have been previously implicated in the pathogenesis of autoimmune disorders including rheumatoid arthritis (free and total IGF1) (24), Crohn’s disease (total IGF1) (25, 26), and T1D (total IGF1) (22). IGF1 has previously been shown to promote the *in vitro* proliferation of sorted human Tregs (27). Additionally, IGF1 treatment inhibited T1D development in the NOD and streptozotocin (STZ)-induced models, with the latter ascribed to increased Treg proliferation in peripheral blood, leading to elevated numbers of Tregs in the pancreas (27). Likewise, the development of murine experimental autoimmune encephalitis (EAE) (27) and allergic contact dermatitis (28) were inhibited by IGF1, with elevated numbers of Foxp3^+^ Tregs observed at the site of autoimmunity, as a direct consequence of Treg-specific IGF1:IGF1 receptor (IGF1R) signaling. These studies suggest that IGF1 may induce immunoregulation through the preferential induction of Treg proliferation as compared to Tconv.

Despite this propensity for IGF1 to impact Tregs, the mechanisms by which this growth factor engages the IGF1R and induces downstream PI3K/Akt signaling (29) and proliferation of Treg from bulk CD4^+^ T cell populations remains unclear. Additionally, IL-2 and IGF1 combination treatment has yet to be evaluated as a synergistic approach toward promoting Treg expansion. Therefore, we hypothesized that IGF1 may augment the extent to which IL-2 preferentially induces Treg expansion. Herein, we demonstrate IL-2 and IGF1 synergism in murine and human Tregs while providing mechanistic insights and establishing evidence for the feasibility of *in vivo* therapeutic and *in vitro* laboratory applications of this combinatorial treatment.

## RESULTS

### IGF1R Expression During T Cell Development and Activation in NOD mice

We hypothesized that IGF1 may preferentially induce Treg proliferation due to increased IGF1R expression on this subset as compared to Tconv. Thus, we measured IGF1R expression by flow cytometry on NOD.Foxp3-GFP thymocyte and splenocyte populations through the stages of central development and peripheral activation, respectively. IGF1R expression was highest on CD4^-^CD8^-^ double negative (DN) thymocytes (geometric mean fluorescence intensity [gMFI] of 472.20 ± 93.30) as compared to later developmental stages (CD4^+^CD8^+^ double positive (DP), 174.20 ± 52.67; CD4^+^CD8^-^ single positive (SP), 271.2 ± 69.76; CD4^-^CD8^+^ SP, 327.20 ± 96.90; **Fig. 1A-B**, **Supplementary Fig. 1A**), in agreement with a previous study showing that human DN thymocytes express approximately three times more surface IGF1R protein than CD4^+^CD8^+^ DP thymocytes (30). IGF1R expression rebounded partially in CD4^+^CD8^-^ and CD4^-^CD8^+^ SP thymocytes (**Fig. 1A-B**, **Supplementary Fig. 1A**), remaining lower than in DN thymocytes. In the periphery, splenic CD4^+^Foxp3^-^CD44^hi^CD62L^-^ memory Tconv showed significantly higher levels of IGF1R (468.80 ± 51.37) than CD4^+^Foxp3^-^ CD44^lo^CD62L^+^ naïve Tconv (209.60 ± 24.63, **Fig. 1C-D**, **Supplementary Fig. 1B**), as previously reported in Balb/c mice (31). This expression pattern was replicated upon comparing naïve and memory CD4^+^Foxp3^+^ Treg (178.22 ± 75.04 vs. 337.80 ± 83.02, **Fig. 1C-D**, **Supplementary Fig. 1B**), suggesting that differential IGF1R levels may partially account for IGF1-mediated T cell regulation via impacts on memory Tregs in mice.

**Figure 1.**
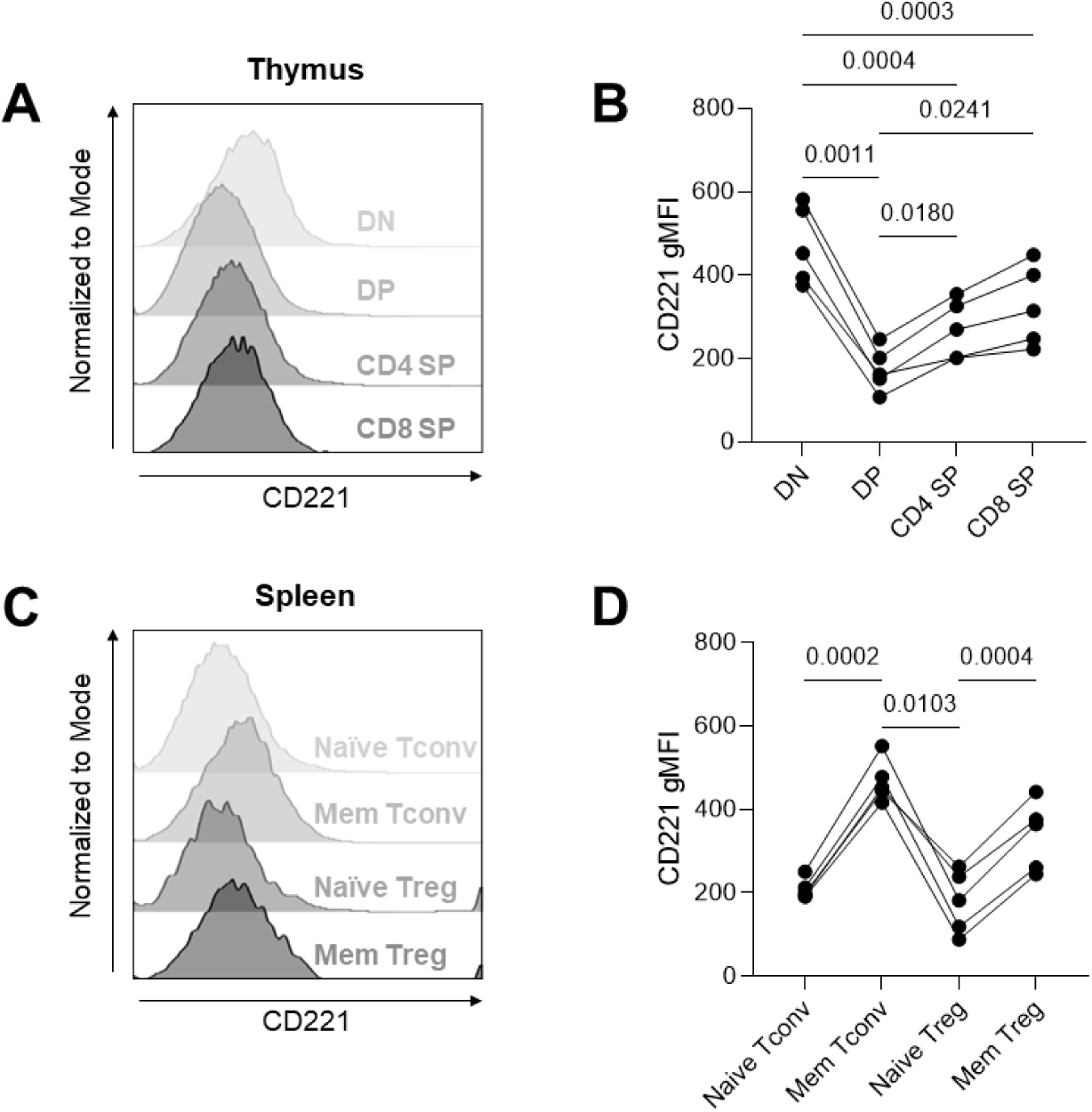
Murine IGF1R expression across the maturation stages of Tconv and Treg. Thymus and spleen of pre-diabetic NOD.Foxp3-GFP mice were stained for IGF1R and differentiation of CD4^+^ and CD8^+^ naïve and memory T cells via flow cytometry. (**A**) Representative histogram of IGF1R/CD221 expression on CD4^-^CD8^-^ double negative (DN), CD4^+^CD8^+^ double positive (DP), CD4^+^CD8^-^ single positive (CD4 SP), and CD4^-^CD8^+^ (CD8 SP) thymocytes. (**B**) Geometric mean fluorescence intensity (gMFI) of IGF1R at various stages of thymocyte development. (**C**) Representative histogram of IGF1R expression on naïve (CD62L^+^CD44^lo^) and memory (CD62L^-^CD44^hi^) Tconv (CD4^+^Foxp3^-^) and Treg (CD4^+^Foxp3^+^) populations in spleen. (**D**) gMFI of IGF1R on CD4^+^ T cell subpopulations. Repeated measures one-way ANOVA with Bonferroni’s multiple comparisons test. n = 5 mice.

### IGF1 and IL-2 Synergize *In Vitro* to Preferentially Induce IGF1R Signaling in NOD Tregs

IGF1R signaling was assessed in splenocytes from NOD mice via Phosflow staining to determine whether IGF1R protein expression directly correlated with the magnitude of IGF1R signaling. Modest IGF1-mediated induction of pS6 (Ser235/236), a measure of PI3K/Akt signaling downstream of IGF1R (32), was observed in CD4^+^Foxp3^+^Helios^+^ Tregs (1.38-fold increase at 15- and 60-minutes post-treatment, **Fig. 2A-B**, **Supplementary Fig. 2**). Thus, we hypothesized that IGF1 could potentially synergize with other inducers of the PI3K/Akt pathway, such as cytokines, to augment Treg IGF1R signaling. In particular, IL-2 has previously been shown to preferentially induce the proliferation of Tregs (33). Therefore, we treated murine splenocytes with low-dose IL-2 in combination with IGF1 in order to test whether IGF1 could further enhance Treg PI3K/Akt signaling. Indeed, the combination enhanced Treg-specific pS6 in a synergistic manner (2.50- and 3.22-fold increase at 15 and 60 minutes, respectively) beyond that of either agent alone (IGF1: 1.38-fold increase at 15- and 60-minutes, IL-2: 1.49- and 1.91-fold increase at 15- and 60-minutes post-treatment, respectively, **Fig. 2A-B**, **Supplementary Fig. 2**). Importantly, the synergy was specific to the Treg compartment, as the combination treatment did not significantly increase pS6 in naïve or memory Tconv (**Fig. 2A-B**, **Supplementary Fig. 2**). In contrast to the PI3K/Akt pathway, IL-2R signaling, as demonstrated by pSTAT5 (Tyr694) induction by IL-2 (**Fig. 2C-D**, **Supplementary Fig. 2**), was unaffected by the addition of IGF1 (**Fig. 2E-F**, **Supplementary Fig. 2**). Together, these data imply that IGF1 and IL-2 are capable of synergizing to promote Treg-specific IGF1R signaling and thereby may promote downstream effects of PI3K/Akt signaling such as cellular proliferation and survival.

**Figure 2.**
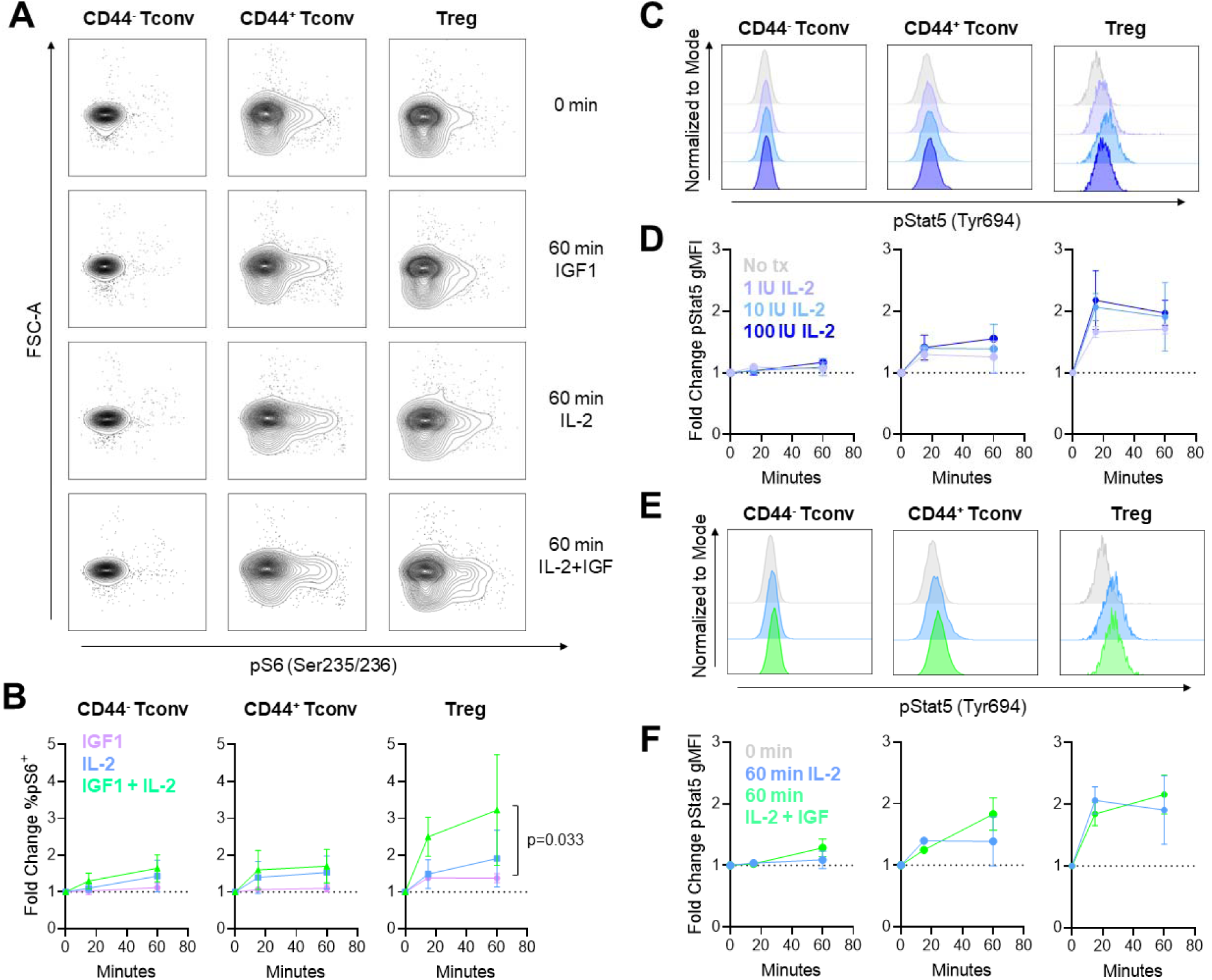
IGF1 synergizes with LD IL-2 to promote PI3K/Akt signaling specifically in Tregs of NOD mice. Splenocytes of pre-diabetic mice (14-22 weeks old) were stimulated with IGF1 (100 ng/mL, n = 3, purple), IL-2 (10 IU/mL, n = 5, blue), or IGF1 + IL-2 (n = 5, green) for 15 or 60 minutes prior to phosflow staining. (**A**) Representative contour plots of pS6 (Ser235/236) induction in CD44^-^ Tconv (CD4^+^Foxp3^-^Helios^-^), CD44^+^ Tconv, and Treg (CD4^+^Foxp3^+^Helios^+^). (**B**) Data were normalized by % pS6^+^ cells without treatment/mouse to calculate fold change. Repeated measures two-way ANOVA with Tukey’s multiple comparisons test. CD44^-^ Tconv: Time x Treatment, p = 0.238; Time, p = 0.004; Treatment, p = 0.091. CD44^+^ Tconv: Time x Treatment, p = 0.234; Time, p = 0.003; Treatment, p = 0.201. Treg: Time x Treatment, p = 0.032; Time, p = 0.003; Treatment, p = 0.033. (**C**) Representative histograms of pStat5 (Tyr694) induction in CD44^-^ Tconv, CD44^+^ Tconv, and Treg without treatment (gray) or treated with 1 IU/mL (periwinkle), 10 IU/mL (aqua), or 100 IU/mL (blue) rhIL-2. n = 2 mice. (**D**) Data in (C) were normalized by pStat5 gMFI without treatment/mouse to calculate fold change. (**E**) Representative histograms of pStat5 (Tyr694) induction in CD44^-^ Tconv, CD44^+^ Tconv, and Treg without treatment (gray) or treated for 60 minutes with 10 IU/mL rhIL-2 (blue) or 10 IU/mL rhIL-2 + 100 ng/mL rhIGF1 (green). n = 2 mice. (**F**) Data in (E) were normalized by pStat5 gMFI without treatment/mouse to calculate fold change.

### IGF1 and IL-2 Synergize *In Vivo* to Promote Treg Proliferation

To assess whether our *in vitro* observation of IGF1 and IL-2 treatment supporting Treg-specific IGF1R signaling would translate to *in vivo* Treg expansion, pre-diabetic NOD mice were treated with 10 μg rhIGF1 twice daily for three weeks (34) and/or IL-2 Antibody Complex (IAC), a method commonly used to extend the half-life of low-dose IL-2, every other day for one week (35) (**Fig. 3A**). We examined populations known to proliferate in response to IL-2 treatment, including CD8^+^ T cells, CD4^+^Foxp3^-^ Tconv, CD4^+^Foxp3^+^ Tregs, and NK cells (18, 19), in the peripheral blood of treated mice. As expected, IAC treatment for one week increased the CD4^+^Foxp3^+^ Treg percentage (**Fig. 3B-C**) with some off-target CD8^+^ T cell and NK cell proliferation (**Supplementary Fig. 3**). While IAC did not significantly increase overall Treg proliferation, the majority of proliferating Tregs occurred in the Helios^+^ thymically-derived (tTreg, 59.61 ± 2.82%) over the Helios^-^ peripherally-induced (pTreg) population (29.10 ± 5.03%) (**Fig. 3D-E**) (36, 37). In contrast, one week of IGF1 treatment resulted in significantly increased Treg proliferation compared to PBS- and IAC-treated mice, with proliferation observed in both the Helios^+^ and Helios^-^ Treg fractions (**Fig. 3D-E**), suggesting that IGF1 can stimulate both tTreg and pTreg *in vivo*. Importantly, the combination of IGF1 plus IAC resulted in a significant increase in overall CD4^+^Foxp3^+^ Treg percentage (12.50 ± 2.57%) compared to IGF1 or IAC alone (5.32 ± 0.77% or 6.83 ± 0.77%, respectively) or PBS control (3.54 ± 0.41) at one week after the start of treatment (**Fig. 3B-C**) without increasing off-target CD8^+^ T cell or NK cell proliferation beyond IAC alone (**Supplementary Fig. 3**). This resulted in a significant fold-increase in CD4^+^Foxp3^+^ Tregs from baseline in the IGF1 + IAC group (3.4 ± 0.42) compared to IAC and IGF1 treatment alone (2.64 ± 0.74 and 1.24 ± 0.19) (**Fig. 3B-C**). While the percentage of CD4^+^Foxp3^+^ Tregs remained increased in the IGF1 + IAC group at two weeks post-treatment as compared to PBS control, the levels were already beginning to contract to those observed pre-treatment and were no longer statistically different from other treatment groups by three weeks post-treatment (**Fig. 3B-C**). These findings suggest that IGF1 and low-dose IL-2 synergize to transiently promote Treg-specific proliferation *in vivo.* These observations then led us to test the question of whether the combination treatment would impact human immune cells similarly.

**Figure 3.**
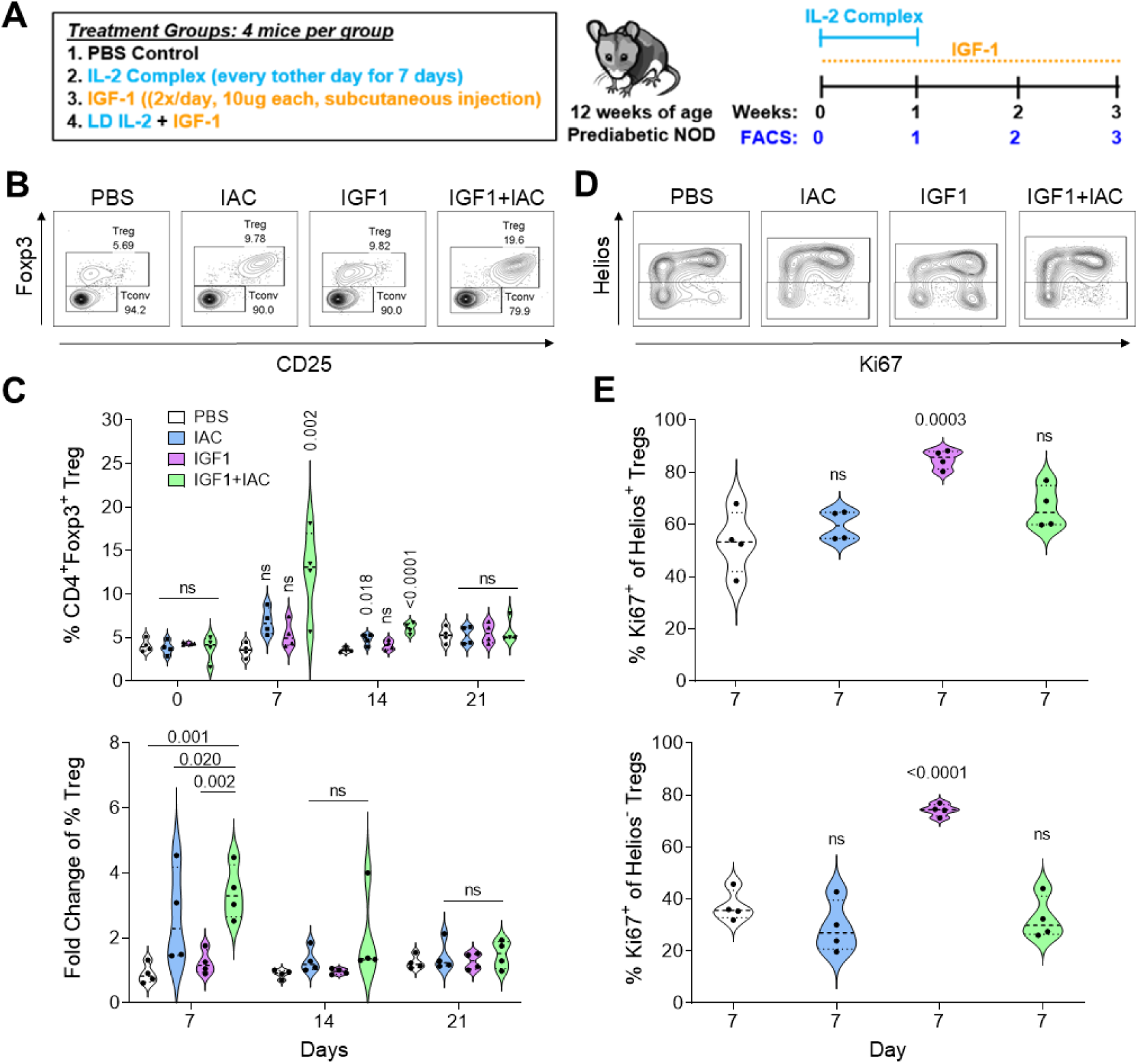
IGF1 treatment stimulates Tregs and synergizes with IL-2. (**A**) Experimental scheme. Prediabetic NOD mice were treated with PBS, IL-2 antibody complex [IAC: anti-IL-2 (JES6.1) + rmIL-2], IGF1, or IGF1 + IAC starting at 12 weeks of age. IL-2 complex was delivered every other day for 7 days, while IGF-1 was delivered as 10 µg subcutaneous injections twice a day for a duration of 3 weeks. Peripheral blood was collected for flow cytometric analysis before treatment and weekly for up to 3 weeks post-treatment. (**B**) Representative dot plots showing CD4^+^Foxp3^+^CD25^+^ Treg and CD4^+^Foxp3^-^ Tconv in peripheral blood after one week of treatment. (**C**) Percentage of CD4^+^Foxp3^+^CD25^+^ Treg at 0, 7, 14, and 21 days in PBS (white), IAC (blue), IGF1 (purple), and IGF1+IAC (green) treated groups. One-way ANOVA with Dunnett’s multiple comparison to the PBS timepoint. Fold change of Treg over day 0 in the peripheral blood 1, 2, and 3 weeks after the start of treatments. One-way ANOVA with Tukey’s multiple comparison at each timepoint. Plots were pre-gated on CD45^+^CD19^-^ Ly6G^-^ cells. (**D**) Representative dot plots showing Helios and Ki67 expression on CD4^+^Foxp3^+^ Tregs in peripheral blood after one week of treatment. (**E**) Percentage of Ki67^+^ cells in Helios^+^ Tregs or Helios^-^ Tregs. One-way ANOVA with Dunnett’s multiple comparison to the PBS timepoint. n = 4 mice per group. ns = not significant.

### Naïve Human Tregs Express High Levels of IGF1R

To understand if differential IGF1R levels observed on T cell subsets from the T1D-prone NOD mouse may translate to human subjects, IGF1R (CD221) expression was quantified by flow cytometry on CD4^+^ T cell subsets from fresh whole blood of children aged 4-16 years with and without T1D (n=14 and n=15, respectively; **Table 1**). Analyzing the total cohort together, IGF1R expression was significantly higher on naïve (CD45RA^+^CD197^+^) versus memory (CD45RA^-^) CD4^+^ T cells (**Fig. 4A-B**, **Supplementary Fig. 4**), in agreement with previous reports from human studies (38) but in direct contrast with findings in mice (**Fig. 1C-D**, **Supplementary Fig. 1B**) (31). Intriguingly, naïve Tregs (CD45RA^+^CD197^+^CD25^hi^CD127^lo/-^) showed significantly higher IGF1R expression than all other subsets assessed, including naïve Tconv (CD45RA^+^CD197^+^CD127^+^) (**Fig. 4A-B**, **Supplementary Fig. 4**), suggesting that naïve Tregs may preferentially respond to IGF1 in humans.

**Figure 4.**
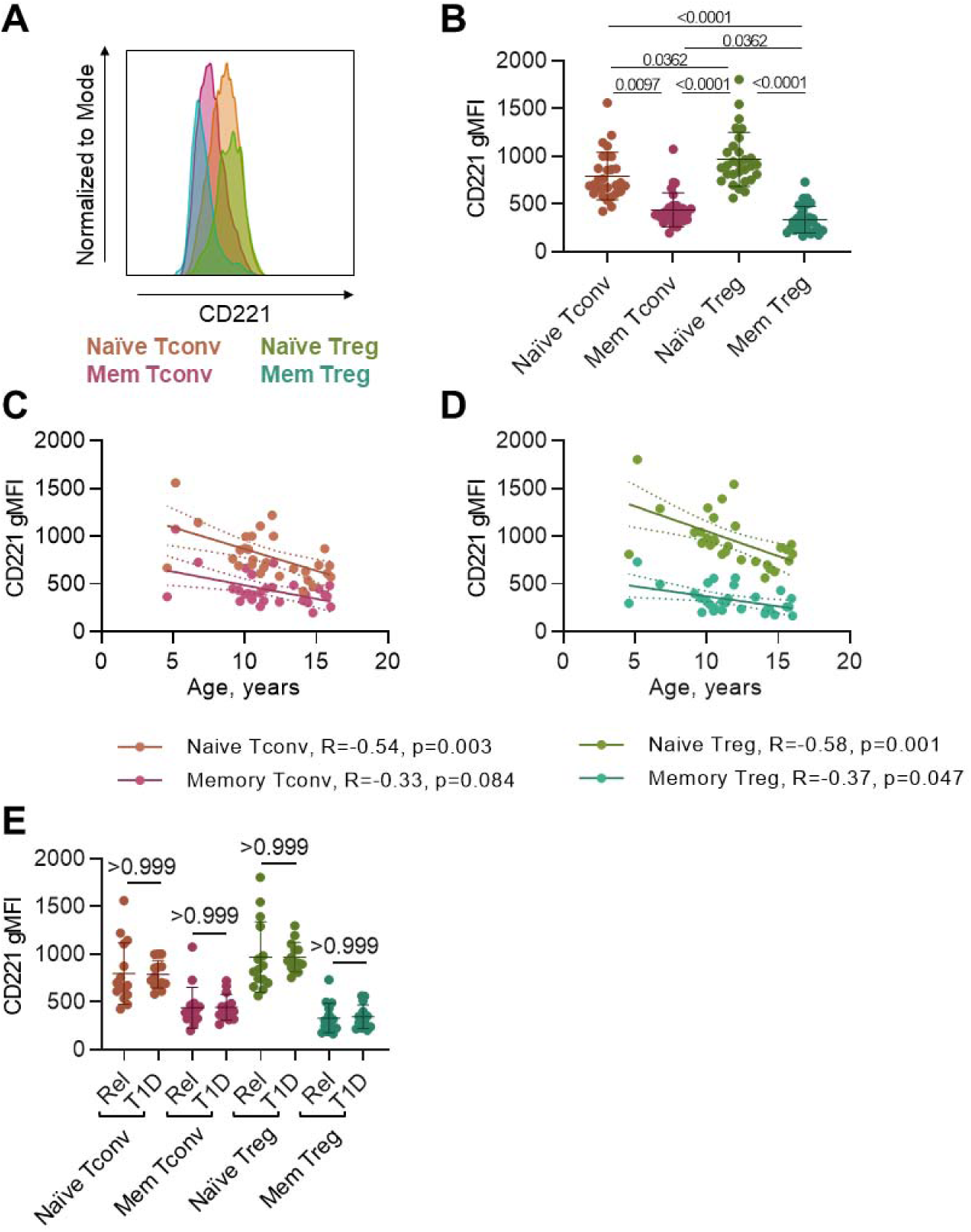
Naive human Treg express significantly higher levels of IGF1R than naïve Tconv. Whole blood staining was performed to further classify CD3^+^CD4^+^ T cells into naïve (CD45RA^+^CD197^+^), memory (CD45RA^-^), conventional (CD127^+^), and regulatory (CD25^hi^CD127^lo/-^) subsets. (**A**) Representative histogram of geometric mean fluorescence intensity (gMFI) of IGF1R (CD221) on naïve Treg (green), naïve Tconv (orange), memory Tconv (pink), and memory Treg (blue) within one subject. (**B**) IGF1R expression is highest on naïve Treg, followed closely by naïve Tconv. Memory Tconv and memory Treg show lowest IGF1R expression. Friedman test with Dunn’s multiple comparisons test. Whole blood staining revealed a negative correlation between IGF1R gMFI and age of subject in years for (**C**) naïve and memory Tconv and (**D**) naïve and memory Treg subsets. Best fit third-order polynomial function shown (solid line, blue) with 95% confidence intervals (dashed lines, blue). Spearman correlation R and p values shown below. (**E**) Comparison of CD221 gMFI on previously defined subsets between T1D and first-degree relatives (Rel) of T1D subjects. Kruskal-Wallis test with Dunn’s multiple comparisons test. n = 29 subjects.

**Table 1.** Demographic information for relatives and T1D subjects enrolled in whole blood IGF1R expression study.

| Cohort | Relatives | T1D |
| --- | --- | --- |
| Total Subjects, n | 15 | 14 |
| Sex, n (%) |  |  |
| Male | 7 (47) | 7 (50) |
| Female | 8 (53) | 7 (50) |
| Age (years) | 12.3 ± 3.2 | 10.9 ± 2.8 |
| Height (m)* | 1.5 ± 0.2 | 1.4 ± 0.2 |
| Weight (kg)* | 45.4 ± 17.5 | 39.1 ± 13.7 |
| BMI (kg/m<sup>2</sup>)* | 18.2 ± 3.7 | 18.8 ± 3.7 |
| Race/Ethnicity, n (%) |  |  |
| Caucasian | 13 (87) | 7 (50) |
| Hispanic | 0 (0) | 3 (21) |
| African-Am | 0 (0) | 1 (7) |
| Asian/Pac-Isl | 1 (7) | 0 (0) |
| Other | 1 (7) | 3 (21) |
| Disease Duration (years) | N/A | 3.0 ± 2.3 |

\* Provision of height and weight was voluntary; thus, these data are available for some, but not all study subjects.

*IGF1R* mRNA expression has previously been shown to decrease in human peripheral blood mononuclear cells (PBMCs) with aging in adult subjects (39); however, immune subset-specific IGF1R expression has been poorly characterized. We observed that IGF1R levels displayed a significant negative correlation with subject age in a pediatric cohort in the naïve Tconv (R = -0.54, p = 0.003, **Fig. 4C**) and naïve Treg (R = -0.58, p = 0.001, **Fig. 4D**) compartments, and this association was weaker but still apparent in memory Tconv (R = -0.33, p = 0.084, **Fig. 4C**) and memory Treg (R = -0.37, p = 0.047, **Fig. 4D**) subsets. Collectively, our findings suggest that IGF1 may preferentially induce signaling in naïve Tregs, particularly in early life when T cell maturation remains active (40).

We also compared IGF1R expression on T cell subsets from subjects with and without T1D (**Table 1**) in order to determine whether disease status might impact the degree of IGF1 signaling in CD4^+^ T cells. In contrast to known modulation of peripheral IGF1 levels in T1D (22), IGF1R levels on naïve Tconv, memory Tconv, naïve Treg, and memory Treg were similar when comparing diabetes-free relatives to age- and sex-matched subjects with T1D (**Fig. 4E**). These data suggest that IGF1R expression on CD4^+^ T cells is not impaired in T1D subjects.

### IGF1 Enhances IL-2-Mediated pSTAT5 Signaling in Naïve Human Tregs

Differences in receptor expression suggest that IGF1 may preferentially induce IGF1R signaling in naïve Treg as compared to other CD4^+^ T cell subsets, although to our knowledge, this question has yet to be formally tested. Therefore, we measured whether IGF1 preferentially augmented phosphorylation of PI3K/Akt pathway targets downstream of IGF1R in human CD4^+^ T cell subsets (**Table 2**). While IGF1 treatment alone was unable to induce pS6 (Ser235/236) expression (**Fig. 5A-B**, **Supplementary Fig. 5**), we surmised that IGF1 could potentially synergize with IL-2 to preferentially induce proliferative signaling downstream of IGF1R in naïve human Tregs, similar to that observed in murine Tregs (**Fig. 2**). However, unlike our findings in mouse, combination treatment of human PBMC with IL-2 + IGF1 did not show a significant impact on pS6 expression in CD4^+^ T cells (**Fig. 5A-B**, **Supplementary Fig. 5**). Surprisingly, IL-2 + IGF1 synergism in the human setting appeared in the IL-2 signaling pathway as measured via pSTAT5 (Tyr694) (**Fig. 5C-D**, **Supplementary Fig. 5**). While pSTAT5 induction in naïve Treg (CD45RA^+^FOXP3^+^Helios^+^) was observed at 15 minutes after treatment with IL-2 as compared to IGF1 alone (gMFI of 14748.50 ± 1792.13 vs. 10747.67 ± 1349.73, p = 0.004), that effect was lost by 60 minutes post-treatment (13518.33 ± 3256.56 vs. 10640.00 ± 1009.38, p = 0.178). In contrast, IL-2 + IGF1 sustained pSTAT5 in naïve Treg to the end of the timecourse in comparison to IGF1 alone (14758.50 ± 1599.26 vs. 10747.67 ± 1349.73, p = 0.002 at 15 minutes, 15619.67 ± 1699.05 vs. 10640.00 ± 1009.38, p = 0.001 at 60 minutes post-treatment, **Fig. 5C-D**, **Supplementary Fig. 5**). A similar pattern of the combination treatment sustaining pSTAT5 signaling past that of IL-2 alone was observed in naïve Tconv (CD45RA^+^FOXP3^-^Helios^-^, IL-2 + IGF1: 3558.33 ± 543.85 vs. IGF1: 2625.00 ± 416.35, p = 0.020; IL-2: 3626.67 ± 529.56 vs. IGF1: 2625.00 ± 416.35, p = 0.012 at 15 minutes, IL-2 + IGF1: 3716.67 ± 501.74 vs. IGF1: 2572.33 ± 645.68, p = 0.018; IL-2: 3371.67 ± 764.61 vs. IGF1: 2572.33 ± 645.68, p = 0.175 at 60 minutes post-treatment, **Fig. 5C-D**), possibly due to low CD25 expression. On the other hand, CD25^hi^ memory Tconv (CD45RA^-^FOXP3^-^Helios^-^) showed sustained pSTAT5 levels not only with the combination treatment but also in the IL-2 alone condition (IL-2 + IGF1: 10205.00 ± 1271.44 vs. IGF1: 3185.17 ± 294.48, p < 0.0001; IL-2: 10279.67 ± 1540.80 vs. IGF1: 3185.17 ± 294.48, p = 0.0002 at 15 minutes, IL-2 + IGF1: 10283.83 ± 1617.34 vs. IGF1: 3080.17 ± 603.34, p < 0.0001; IL-2: 8801.17 ± 3519.48 vs. IGF1: 3080.17 ± 603.34, p = 0.023 at 60 minutes post-treatment, **Fig. 5C-D**). Together, these findings suggest that IL-2 + IGF1 preferentially augments IL-2R signaling in the presence of IGF1R signaling in naïve human CD4^+^ T cells.

**Figure 5.**
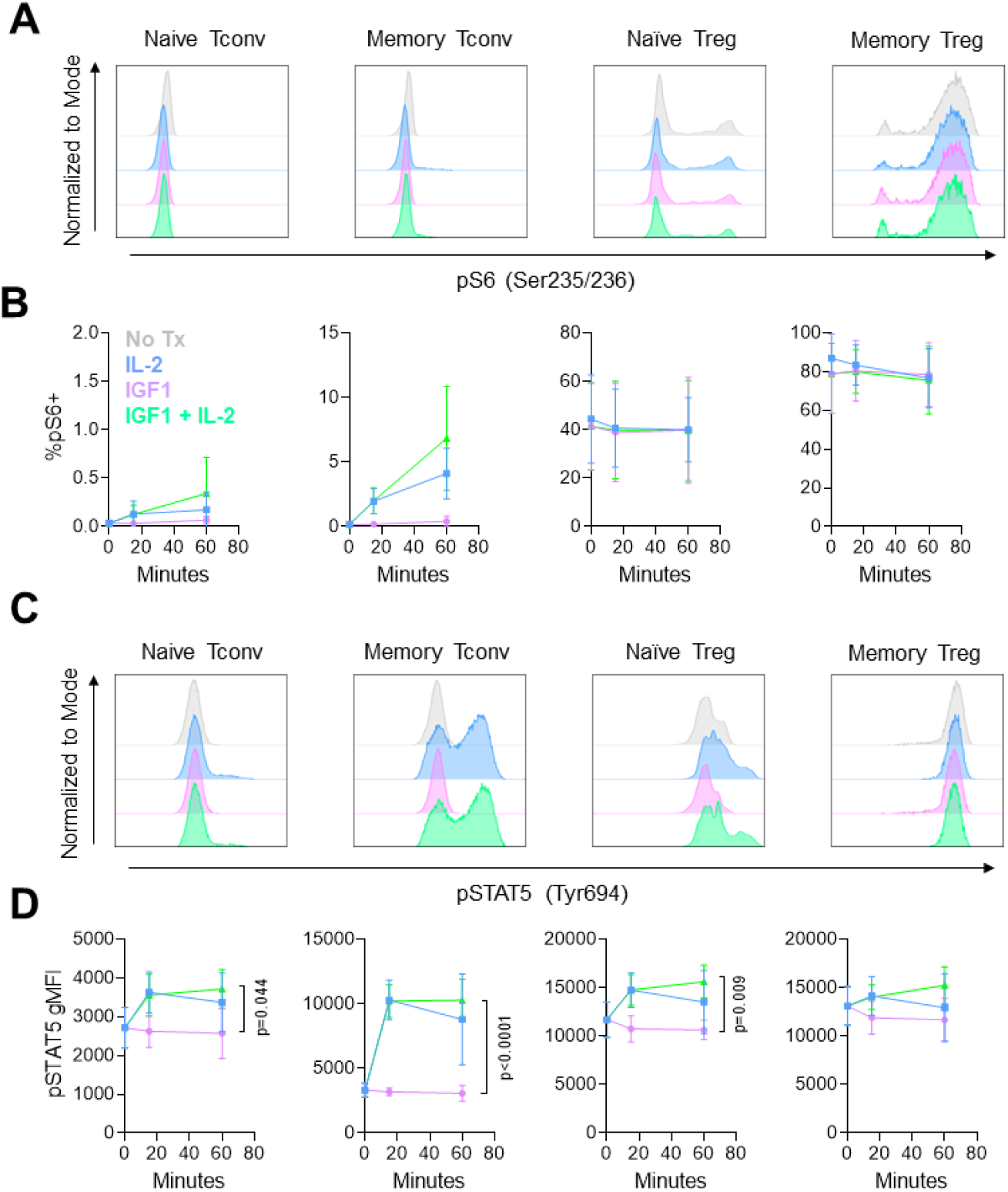
IGF1 enhances IL-2-mediated pSTAT5 signaling in naïve human Tregs. PBMC were stimulated with IL-2 (20 IU/mL, blue), IGF1 (100 ng/mL, purple), IGF1 + IL-2 (green), or no treatment (No Tx, gray) for 15 or 60 minutes prior to phosflow staining. (**A**) Representative histograms of pS6 (Ser235/236) induction in naive Tconv (CD4^+^CD45RA^+^Foxp3^-^Helios^-^), memory Tconv (CD4^+^CD45RA^-^Foxp3^-^Helios^-^), naïve Treg (CD4^+^CD45RA^+^Foxp3^+^Helios^+^), and memory Treg (CD4^+^CD45RA^-^Foxp3^+^Helios^+^). (**B**) Quantification of percentage of pS6^+^ cells over the timecourse. (**C**) Representative histograms of pSTAT5 (Tyr694) induction. (**D**) Quantification of pSTAT5 gMFI over the timecourse. Repeated measures two-way ANOVA with Tukey’s multiple comparisons test. Naïve Tconv: Time x Treatment, p = 0.003; Time, p = 0.001; Treatment, p = 0.045. Memory Tconv: Time x Treatment, p < 0.0001; Time, p < 0.0001; Treatment, p < 0.0001. Naïve Treg: Time x Treatment, p = 0.0002; Time, p = 0.002; Treatment, p = 0.012. Memory Treg: Time x Treatment, p = 0.060; Time, p = 0.839; Treatment, p = 0.187. n = 6 subjects.

**Table 2.** Demographic information for subjects enrolled in IGF1R signaling, proliferation, receptor subunit regulation, and gene editing studies.

| Cohort | Age | Sex |
| --- | --- | --- |
| *IGF1R Signaling* | 18 | F |
|  | 19 | M |
|  | 19 | F |
|  | 24 | M |
|  | 24 | M |
|  | 24 | F |
| *Homeostatic Proliferation* | 16 | F |
|  | 18 | M |
|  | 26 | F |
|  | 41 | F |
|  | 43 | F |
| *Receptor Subunit Regulation* | 18 | M |
|  | 18 | F |
|  | 19 | F |
|  | 20 | F |
|  | 21 | M |
|  | 22 | M |
|  | 24 | M |
| *Gene Editing* | 18 | M |
|  | 20 | F |
|  | 21 | F |
|  | 22 | F |
|  | 24 | F |

### IGF1 Augments IL-2-Mediated Homeostatic Proliferation of Naïve Human Treg

Our observation that IL-2 + IGF1 treatment augments IL-2R signaling in naïve Treg implied that IGF1 could potentially synergize with IL-2 to preferentially induce the homeostatic proliferation of naïve human Tregs in the absence of T cell receptor (TCR) stimulation (signal 1) or costimulation (signal 2). Thus, we treated bulk naïve CD4^+^ T cells with IL-2 and/or IGF1 to assess Treg proliferation (**Fig. 6A**, **Table 2**). While IGF1 alone was unable to induce the proliferation of naïve FOXP3^+^Helios^+^ Treg (**Fig. 6B**), the addition of IL-2 promoted the expansion of naïve Tregs as compared to IL-2 alone (division index [DI] of 0.45 ± 0.21 vs. 0.08 ± 0.11, **Fig. 6B-C**). Indeed, supplementation of low-dose IL-2 with IGF1 significantly increased the percentage of FOXP3^+^Helios^+^ Tregs within bulk naïve CD4^+^ T cell culture (4.74 ± 2.24 % vs. 3.09 ± 2.18 %, **Fig. 6D-E**). Thus, IGF1 can synergize with low-dose IL-2 to specifically drive the homeostatic proliferation of naïve Treg.

**Figure 6.**
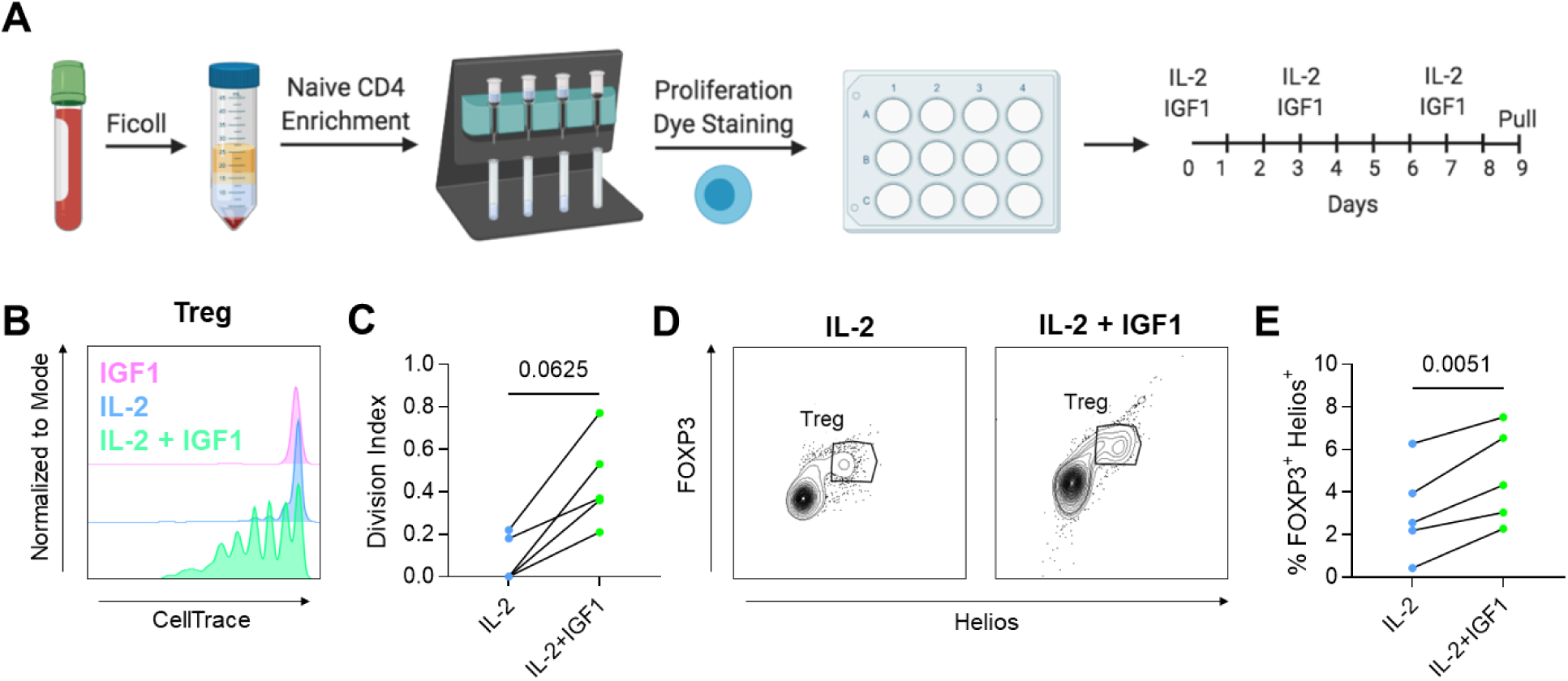
IGF1 augments the IL-2-mediated homeostatic proliferation of naïve Treg. (**A**) Methods for *in vitro* naïve CD4^+^ T cell homeostatic proliferation experiments. Density gradient centrifugation was performed to isolate PBMCs from whole blood, followed by magnetic bead-based naïve CD4^+^ T cell enrichment. Cells were stained with Cell Proliferation Dye eFluor670 to track proliferation prior to plating in complete RPMI supplemented with 20 IU/mL IL-2 and/or 100 ng/mL IGF1. Cytokines were replenished on days 3 and 7 and cultures were stained for flow cytometric analysis on day 9-11. Image created with BioRender. (**B**) Representative plots of cell proliferation dye from IGF1 (purple), IL-2 (blue), and IL-2 + IGF1 (green) conditions show (**C**) enhanced proliferation of naïve Treg in the presence of IL-2 + IGF1 versus IL-2 alone, as quantified by the division index. Wilcoxon-test. (**D**) Representative plot of FOXP3 and Helios expression on CD4^+^CD45RA^+^ T cells showing that (**E**) percentage of FOXP3^+^Helios^+^ cells was increased upon treatment with IL-2 + IGF1 as compared to IL-2 alone. Paired t test. n = 5 subjects.

### IGF1 Synergizes with IL-2 to Modulate IL-2R Subunit Expression in Naïve Treg and Tconv

To elucidate the mechanism of IL-2 + IGF1 synergism, we assessed whether IGF1 modulates expression of any of the IL-2R subunits (IL-2Rα, IL-2Rβ, and IL-2Rγ/common γ-chain) or whether IL-2 modulates IGF1R expression by naïve human CD4^+^ T cells (**Table 2**). Upon formation of the IL-2: high-affinity IL-2R complex consisting of all three IL-2R subunits, the complex is known to be internalized, with IL-2Rβ and IL-2Rγ degraded and IL-2Rα recycled to the surface for further use by Tregs (41). Previous murine studies have suggested that IL-2 upregulates IL-2Rα/CD25 expression on Tregs in a positive feedback loop (42), supporting the notion of IL-2Rα recycling guiding high-affinity IL-2R availability. In contrast, IGF1 treatment of human T cells has been shown to promote the endocytosis and degradation of IGF1R, levels of which were dampened for a few days, without evidence of the receptor being recycled (43, 44). Therefore, we hypothesized that IL-2 + IGF1-mediated proliferation of naïve human Tregs might occur as a consequence of CD25 induction. Indeed, we observed that a short-term culture with the combination treatment of IL-2 and IGF1 significantly enhanced CD25 expression on CD45RA^+^CCR7^+^FOXP3^+^Helios^+^ naïve Tregs as compared to treatment with IGF1 alone (gMFI of 36027.57 ± 10817.01 vs. 20246.57 ± 9472.10, **Fig. 7A-B**, **Supplementary Fig. 6**). While we did not observe evidence that IL-2 + IGF1 treatment regulates levels of the intermediate-affinity IL-2R subunits, CD122 and CD132, on the naïve Treg compartment (**Fig. 7A-B**), interestingly, both of these IL-2R subunits were upregulated by IL-2 + IGF1 treatment in CD45RA^+^CCR7^+^FOXP3^-^Helios^-^ naïve Tconv (**Fig. 7C-D**). IL-2Rβ/CD122 levels were significantly increased in the combination treatment and even in the IGF1 alone condition versus IL-2 alone (982.14 ± 64.69 and 965.71 ± 63.41 vs. 828.86 ± 57.46, **Fig. 7C-D**), potentially fine-tuning response to IL-2. Additionally, increased IL-2Rγ/CD132 expression in the IL-2 + IGF1 as compared to no treatment condition (2722.86 ± 278.48 vs. 1760.14 ± 282.61, **Fig. 7C-D**) could possibly allow for broader synergism between IGF1 and other common γ-chain cytokines. IGF1R levels were significantly reduced by IGF1 treatment (**Fig. 7A-D**), as previously reported (43, 44). Together, these findings suggest that IL-2 + IGF1 promotes IL-2 sensitivity in naïve Tregs through induction of CD25 expression and additionally, raises the interesting possibility that IGF1 may enhance sensitivity to γ-chain cytokines in naïve Tconv.

**Figure 7.**
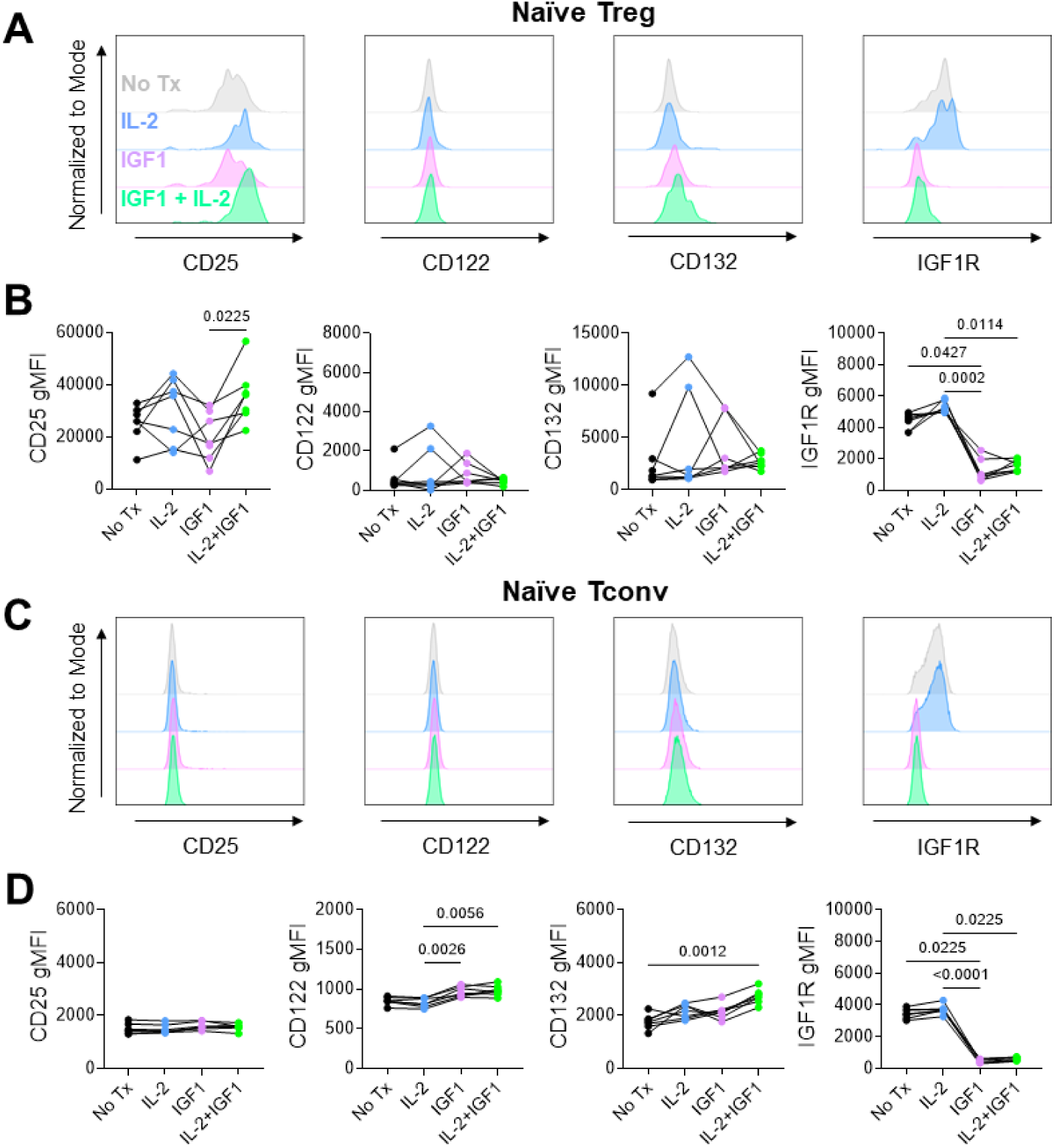
IGF1 synergizes with IL-2 to modulate IL-2R subunit expression in naïve Treg and Tconv. Human PBMC were stimulated with 20 IU/mL rhIL-2 (blue), 100 ng/mL rhIGF1 (purple), IL-2 + IGF1 (green), or neither (No Tx, gray) for two days followed by flow cytometric staining for IL-2R subunits and IGF1R. (**A**) Representative histograms of CD25/IL-2Rα, CD122/IL-2Rβ, CD132/IL-2Rγ, and CD221/IGF1R expression on naïve Treg (CD45RA^+^CD197^+^FOXP3^+^Helios^+^). (**B**) gMFI of each marker on naïve Treg shown per subject across treatment conditions. (**C**) Representative histograms of CD25/IL-2R?, CD122/IL-2Rβ, CD132/IL-2Rγ, and CD221/IGF1R expression on naïve Tconv (CD45RA^+^CD197^+^FOXP3^-^ Helios^-^). (**D**) gMFI of each marker on naïve Tconv shown per subject across treatment conditions. Friedman test with Dunn’s multiple comparisons test. n = 7 subjects.

### IGF1 and **γ**-chain Cytokines Promote CD4^+^ T Cell Transduction While Maintaining Naivety

Maintenance of T cell naivety may prove beneficial in certain translational uses such as quiescent T cell expansion for autologous adoptive cellular therapies (45). Additionally, the creation of antigen-specific T cell therapies for cancer or autoimmunity and the study of rare antigen-specific primary Treg and Tconv responses have been classically hindered by the need for activation to permit transduction with a receptor of interest (46, 47). Thus, we hypothesized that combinatorial IL-2 + IGF1 or IL-7 + IGF1 treatment would enhance transduction of Treg or Tconv, respectively, while avoiding acquisition of a memory phenotype. Indeed, pre-treatment of naïve CD4^+^ T cells with IL-2 + IGF1, followed by transduction with a lentivirally-encoded TCR recognizing the T1D-associated epitope glutamic acid decarboxylase 65 (GAD65) 555-567 (48), significantly enhanced the proliferation of GAD65-reactive TRBV5-1^+^ transduced naïve Tregs beyond IL-2 alone (DI of 1.38 ± 0.91 vs. 0.74 ± 0.69, **Fig. 8A-B**, **Table 2**, **Supplementary Fig. 7**). Similarly, IL-7 + IGF1 treatment significantly increased the DI of TRBV5-1^+^ naïve Tconv as compared to IL-7 treatment alone (1.09 ± 0.52 vs. 0.78 ± 0.55, **Fig. 8A-B**). Despite this increase in proliferation, the overall transduction rate remained similar in IL-2 + IGF1 vs. IL-2 and IL-7 + IGF1 vs. IL-7 conditions (**Fig. 8C-D**). Importantly, Treg and Tconv expressing the lentivirally-encoded T1D-associated TCR remained almost entirely CD45RA^+^CD197^+^ naïve (**Fig. 8E-F**) as opposed to traditional methods that generate mostly memory Treg (49, 50). The resulting naïve antigen-specific Treg and Tconv could then be utilized to study primary activation events, of particular interest in conditions such as T1D wherein many genetic risk loci are tagged to regulators of T cell activation (51, 52).

**Figure 8.**
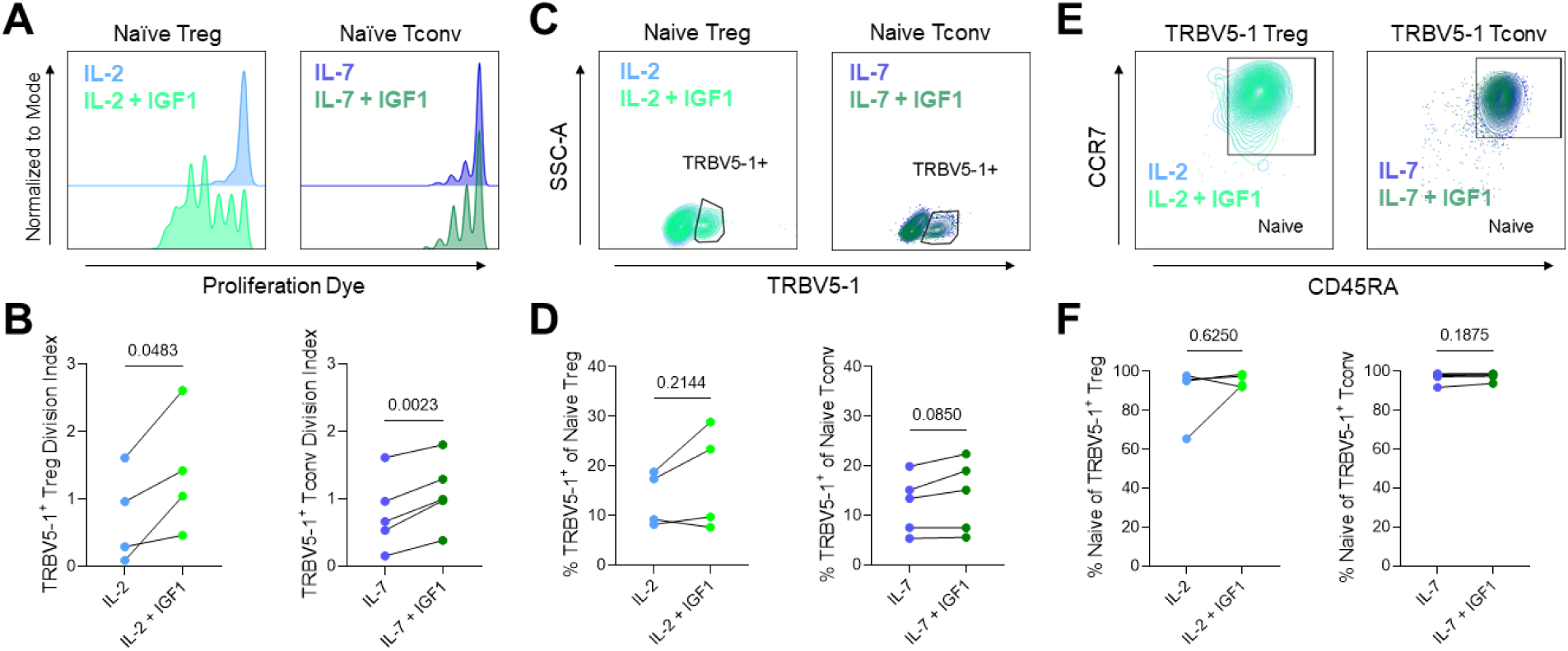
IGF1 and γ-chain Cytokines Promote CD4^+^ T Cell Transduction While Maintaining Naivety. Naïve CD4^+^ T cells were isolated and cultured with 20 IU/mL IL-2 or 10 ng/mL IL-7, with or without 100 ng/mL IGF1, followed by transduction with lentiviral constructs containing T1D-related β-cell-specific T cell receptor (TCR) R164, recognizing glutamic acid decarboxylase (GAD) 555-567 in the context of human leukocyte antigen (HLA)-DRB1*04:01. (**A**) Dye dilution assay shows enhanced proliferation of naïve transduced Treg (CD45RA^+^CD197^+^TRBV5-1^+^FOXP3^+^Helios^+^) with IL-2 + IGF1 (green) as compared to IL-2 (blue) alone and enhanced proliferation of naïve transduced Tconv (CD45RA^+^CD197^+^TRBV5-1^+^FOXP3^-^Helios^-^) with IL-7 + IGF1 (dark green) versus IL-7 alone (dark blue), (**B**) as quantified by division index (DI). Paired t test. (**C**) Representative contour plots of TRBV5-1 expression show similar transduction rates of naïve Treg with IL-2 versus IL-2 + IGF1 and naïve Tconv with IL-7 versus IL-7 + IGF1 treatment. (**D**) Percentage of TRBV5-1^+^ cells transduced with R164 construct. Wilcoxon test. (**E**) Representative contour plots of CD45RA and CD197 expression on TRBV5-1^+^ Treg and Tconv show transduced cells retain naivety. (**F**) Percentage of naïve (CD45RA^+^CD197^+^) cells within the TRBV5-1^+^ Treg and Tconv gates for IL-2 ± IGF1 and IL-7 ± IGF1 treatment, respectively. Paired t test. IL-2 ± IGF1, n = 4 subjects; IL-7 ± IGF1, n = 5 subjects.

## DISCUSSION

While IL-2 is widely-appreciated as an important regulator of Treg expansion and functional maintenance, the potential for off-target effects begs the question of whether adjunctive treatments may synergize with and enable the use of low IL-2 doses. Here, we sought to evaluate whether IGF1 augmented IL-2-mediated Treg expansion, initially characterizing patterns of IGF1R expression on T cell subsets. In agreement with our findings, previous studies support the notion that naïve human CD4^+^ T cells express significantly higher levels of IGF1R than memory CD4^+^ T cells (38, 53), while this pattern is reversed in mice (31, 54). Despite this incongruity, we found novel evidence of naïve human Tregs expressing significantly elevated levels of IGF1R as compared to naïve Tconv. This implies a potential role for IGF1 in maintaining the naïve Treg pool. When taken together with our findings that younger human subject age and earlier murine thymocyte developmental stage are associated with increased IGF1R expression, these data suggest that IGF1 signaling may be important for naïve Tregs in early development, with potential implications for modulating the survival or proliferation of Tregs at immature thymocyte stages (55).

While IGF1 alone had a negligible impact on IGF1R signaling and Treg expansion, we report the novel observation of IGF1 synergizing with IL-2 to enhance proliferative signaling in the absence of TCR ligation. IL-2 + IGF1 preferentially induced activation of the PI3K/Akt pathway in murine Tregs, while the combination treatment instead enforced the IL-2R signaling pathway in CD4^+^ T cells in our human studies. This is in direct contrast to previous studies which have demonstrated induction of PI3K/Akt signaling upon IGF1 treatment of bulk primary human T cells (29, 56). However, these studies involved pre-activation of T cells via TCR agonists (29, 56), suggesting that IGF1 may act as a regulator to amplify signaling pathways already engaged by T cells. Additionally, in agreement with our findings of pSTAT5 induction by the combination treatment in human cells, the literature suggests IGF1R may enhance STAT5 signaling, albeit in different cell types than studied here (57, 58), which could augment well-described IL-2-mediated STAT5 induction (18, 59). Others have previously shown *in vitro* evidence of pS6 induction in murine Tregs by a truncated analog of IGF1 (32) in addition to modest increases in pS6 by IL-2 (60), supporting our observation of IL-2 + IGF1 synergism in promoting PI3K/Akt signaling in murine Tregs. We suspect that the differing observations between mouse and humans may be accounted for by high basal levels of pS6 in human Tregs. Additional experiments with extended timepoints could help to further elucidate reliance on JAK-STAT5 versus PI3K/Akt signaling pathways as proliferative status dramatically remodels cellular signaling and metabolism (61).

Additionally, our studies show that IGF1 can specifically augment human tTreg proliferation *in vitro* and murine pTreg and tTreg *in vivo*, in combination with low-dose IL-2. Others have previously shown that IGF1 alone can induce Treg skewing in human PBMC (62, 63), analogous to pTreg induction observed here in NOD mice. However, our work is unique in that it establishes the synergism of IGF1 with IL-2 and highlights the specificity of this treatment regimen for induction of naïve human Treg proliferation as opposed to a memory Treg phenotype in mouse. Furthermore, we observed that IL-2 + IGF1 upregulated expression of the high-affinity IL-2R subunit, CD25, in naïve human Tregs to promote sensitivity to IL-2, in agreement with our findings of combination treatment enhancing pSTAT5 induction. While culture with cellular supernatants containing IGF1 has been shown to increase CD25 expression by induced human Tregs (iTreg) (62), which are memory Treg due to the requirement for TCR activation for iTreg skewing (36), our study is the first to corroborate similar findings in naïve tTregs.

Beyond enhancing CD25 expression on naïve Tregs, we observed that IGF1 +/- IL-2 upregulated expression of intermediate-affinity IL-2R subunits, CD122 and CD132, on naïve human Tconv. Indeed, a previous study suggested that inactivation of FoxO1, a signaling mediator downstream of IGF1R whose activity is inhibited by IGF1 (29), upregulates expression of CD122 in naïve murine CD4^+^ T cells (64). There is a scarcity of literature on modulation of expression of CD132 levels on T cells (65, 66), suggesting that IL-2 + IGF1 may constitute a novel means to sensitize naïve Tconv to cytokines whose receptors utilize the common γ-chain, including but not limited to IL-2 and IL-7 (10). These findings further promote the notion that the influence of IGF1 on the inflammatory/regulatory balance may be highly dependent on the cytokine milieu.

Our observations of IGF1 potentially sensitizing naïve Treg and naïve Tconv to common γ-chain cytokine signaling inspired investigation into the application of IGF1 toward *ex vivo* homeostatic proliferation for gene transfer in primary human T cells. Others have previously shown successful transduction and maintenance of naivety in primary human CD4^+^ and CD8^+^ T cells by pre-treating with IL-2 (67, 68) or IL-7 (67–69), without the use of conventional TCR activation stimuli, but this approach provides low cell yield, limiting their use in downstream experiments. Here, we showed that the addition of IGF1 to IL-2 treatment not only enhanced the proliferation of transduced cells, but that this appeared to preferentially promote expansion of naïve transduced Tregs. We also found that IGF1 augmented IL-7-mediated proliferation of transduced naïve CD4^+^ Tconv, likely to be partially explained by IL-7 signaling through CD132/IL-2Rγ (10), which we found was upregulated on naïve Tconv in the presence of IGF1 and low-dose IL-2. Although this IL-7-mediated proliferation experiment was performed in the absence of rhIL-2, we suppose that cycling Tconv may have produced low levels of IL-2 for autocrine use (9), allowing for CD132/IL-2Rγ upregulation. In addition to permitting the study of primary antigen-specific Treg and Tconv responses, of importance for *in vitro* modeling of autoimmunity and cancer (70), our combination treatment approach also has relevant clinical implications for autologous adoptive cellular therapies (45). Studies in the delivery of chimeric antigen receptor (CAR) T cells in cancer (71, 72) and CAR Tregs in inflammatory models (73) have clearly shown that therapeutic success is reliant on dampening of memory acquisition and hindering the development of cellular exhaustion. CAR Tregs, while proposed for treatment of autoimmunity, likewise require methodological fine-tuning for prevention of exhaustion and loss of lineage stability (74). Future studies should investigate whether the incorporation of IL-2 + IGF1 or IL-7 + IGF1 into CAR Treg or Tconv expansion protocols, respectively, prevents exhaustion to allow for longer duration of efficacy.

Our published work showing significantly lower peripheral IGF1 levels in pre- and post-T1D (22) suggests that IGF1:IGF1R signaling may be impaired during disease development. While IGF1R levels on CD4^+^ T cells were comparable between healthy subjects and those with T1D, further studies of IGF1R expression in pre-T1D may be necessary to rule out IGF1R dysregulation on CD4^+^ T cells throughout the course of T1D pathogenesis. In combination with our observations here, decreased IGF1 signaling may contribute to autoimmunity (23) by inhibiting naïve tTreg proliferation. However, an important caveat to this idea is that other groups have reported increased naïve Tregs in the peripheral blood of children with T1D (75, 76), which could potentially reflect a decrease in this population at the site of autoimmunity. In line with this notion, targeted delivery methods for IL-2 + IGF1 treatment toward the pancreas and/or pLN may need to be considered to rectify known Treg deficits in the pLN in T1D (6, 77). Clearly, future efforts moving beyond peripheral blood to characterize immune phenotypes at the site of autoimmunity will be necessary to test these ideas, which remains a challenge in cases such as this where human and murine biology diverge.

In sum, we report that IGF1 augments naïve human CD4^+^ T cell sensitivity toward common γ-chain cytokines, and that this can be exploited via IGF1 + low-dose IL-2 treatment to drive the homeostatic proliferation of naïve Tregs. We also present differences in murine and human IGF1R expression and signaling, highlighting the need for future studies of this pathway in human cells specifically. Collectively, our findings present a novel strategy toward rectifying the Treg:Tconv imbalance contributing to many autoimmune conditions. These data support additional studies to assess whether IGF1 can increase the IL-2-mediated homeostatic proliferation of naïve Tregs *in vivo* or of autologous engineered Tregs *ex vivo* for translational benefit, representing potential strategies for clinical intervention to prevent or halt the progression of autoimmune disease, such as T1D.

## METHODS

### Murine IGF1R Staining

Spleen and thymus of six-week-old NOD.Foxp3-GFP/cre (Stock No: 008694, Jackson Laboratory) were processed using frosted glass slides and passed through a 40-micron filter to create single cell suspensions. Red blood cells (RBC) were lysed with ammonium-chloride-potassium buffer prior to staining for flow cytometry. Samples were stained with Fixable Live/Dead Near IR (Invitrogen), washed once with stain buffer (PBS + 2% FBS + 0.05% NaN_3_), and Fc receptors were blocked with anti-CD16/32 for 5 minutes at 4°C (Clone 2.4G2, BD Biosciences). Samples were stained for 30 minutes at 4°C with the following anti-mouse antibodies: CD3e-Brilliant Violet (BV) 605 (145-2C11, BioLegend), CD4-PerCP/Cy5.5 (RM4-5, Thermo Fisher Scientific), CD8a-BV711 (53-6.7, BioLegend), CD44-PE-Cy7 (IM7, BioLegend), CD62L-APC (MEL-14, BioLegend), and CD221-PE (3B7, Santa Cruz Biotechnology). Samples were washed once with stain buffer prior to data acquisition on an LSRFortessa (BD Biosciences) and analysis with FlowJo (v10.6.1; Tree Star).

### Murine IGF1R Signaling

Single cell suspensions of splenocytes were generated from 11–17-week-old pre-diabetic NOD/ShiLtJ mice (Stock No: 001976, Jackson Laboratory), followed by RBC lysis. Cells were resuspended at 10^6^/mL in complete RPMI 1640 without L-glutamine (Corning; cRPMI [supplemented with 2 mM GlutaMAX (Gibco), 100 IU and 100 μg/mL each of penicillin-streptomycin solution (Corning), 1 mM sodium pyruvate (Corning), 0.1 mM non-essential amino acids (Gibco), 10 mM HEPES (Gibco), 10% fetal bovine serum (GenClone), and 0.0004% 2-mercaptoethanol (Sigma-Aldrich)]) and treated with 10 IU/mL recombinant human (rh)-IL-2 (Teceleukin) and/or 100 ng/mL rhIGF1 (BioVision) for 15 or 60 minutes at 37°C. Samples were fixed with an equal volume of Cytofix fixation buffer (BD Biosciences) for 10 minutes at 37°C. Live/Dead fixable near-IR dead cell stain kit (Invitrogen) was applied according to manufacturer’s instructions for dead cell exclusion. Cells were washed once with stain buffer, permeabilized with Phosflow perm buffer III (BD Biosciences) for 30 minutes at 4°C, washed twice with stain buffer, and then incubated with anti-CD16/32 (Clone 2.4G2, BD Biosciences) for 5 minutes at 23°C. Samples were stained with the following fluorescently-labeled anti-mouse antibodies at 23°C for 45 minutes: CD3-PerCP-Cy5.5 (17A2), CD4-BV711 (RM4-5), CD44-PE-Cy7 (IM7), Helios-Pacific Blue (22F6) (BioLegend), Foxp3-AlexaFluor (AF)-AF488 (FJK-16s, eBioscience), pS6 Ser235/236-AF647 (D57.2.2E, Cell Signaling Technology), and pSTAT5 Tyr694-PE (47/STAT5, BD Biosciences). Samples were washed once with stain buffer prior to data acquisition on an LSRFortessa (BD Biosciences) and analysis with FlowJo software (v10.6.1; Tree Star).

### Murine IGF1 + Low-Dose IL-2 Treatment

Pre-diabetic 12-week old female NOD/ShiLtJ mice were treated with IL-2 antibody complex [IAC: 5 μg anti-IL-2 (JES6-1A12, eBioscience) + 1 μg recombinant mouse IL-2] injected intraperitoneally every other day for a week as we previously reported (35), 10 μg rhIGF1 injected subcutaneously twice a day for three weeks (34), IAC + IGF1 in combination, or vehicle control (PBS). Peripheral blood was collected before treatment and on a weekly basis during the treatment regimen for flow cytometric analysis. RBC were lysed prior to staining with Live/Dead fixable near-IR dead cell stain kit (Invitrogen). Cells were stained with the following antibodies: CD19-APC-Cy7 (1D3, BD Biosciences), Ly6G-APC-Cy7 (1A8, BD Biosciences), CD122-Biotin (TM-β1, BD Biosciences), CD45-BV510 (30-F11, BD Biosciences), CD5-BV650 (53-7.3, BD Biosciences), CD4-PE-Cy7 (RM4-5, BioLegend), CD8-AF700 (53-6.7, BioLegend), CD25-PE-CF594 (PC61, BD Biosciences), CD335-PE (29A1.4, BD Biosciences), cKIT (CD117)-Brilliant Blue (BB)-700 (2B8, BD Biosciences), CD44-BV605 (IM7, BD Biosciences), prior to a wash with stain buffer and application of a Streptavidin-BV711 (BD Biosciences) conjugate. Samples were fixed and permeabilized with Foxp3 transcription factor staining buffer set according to manufacturer’s protocol (ThermoFisher) before staining with anti-mouse Helios-APC (22F6, BioLegend), Foxp3-eFluor 450 (FJK-16s, ThermoFisher), Eomes-AF488 (Dan11ma, ThermoFisher), and Ki67-BV786 (B56, BD Biosciences). Data were acquired on an LSRII (BD Biosciences) and analyzed with KALUZA software (v2.1).

### Human Subject Enrollment

Study subjects were recruited from the Children with Diabetes-sponsored Friends for Life (FFL) Conference held annually in Orlando, Florida; the general population; or outpatient clinics at the University of Florida (UF; Gainesville, FL); Nemours Children’s Hospital (Orlando, FL); and Emory University (Atlanta, GA). After providing written informed consent, peripheral blood samples were collected into the UF Diabetes Institute (UFDI) Study Bank from non-fasted subjects by venipuncture in heparin-coated vacutainer tubes (BD Biosciences) in accordance with institutional review board (IRB) approved protocols. Leukapheresis-processed blood of healthy donors was also purchased from LifeSouth Community Blood Centers (Gainesville, FL). Samples were selected from relatives of T1D subjects as well as age- and sex-matched T1D patients for fresh whole blood staining, with detailed deidentified demographic information presented in **Table 1**. For the remainder of experiments performed, peripheral blood samples were selected from healthy subjects (**Table 2**).

### Human Whole Blood Flow Cytometry

To determine IGF1R expression, 200 μL of whole blood was stained with the following fluorescently-labeled anti-human antibodies for thirty minutes at 23°C in the dark: CD25-AF488 (BC96), CD127-BV421 (A019D5), CD197-AF647 (G043H7), CD45RA-PerCP-Cy5.5 (HI100), CD3-APC-Fire750 (SK7), CD221-PE (1H7) (BioLegend), CD4-PE-Cy7 (RPA-T4, eBioscience). RBCs were lysed for five minutes at 23°C with 1-step Fix/Lyse Solution (eBioscience), followed by three washes with stain buffer. Data were acquired on an LSRFortessa (BD Biosciences) and analyzed with FlowJo software (v10.6.1; Tree Star).

### Human IGF1R Signaling

Whole blood was processed to peripheral blood mononuclear cells (PBMCs) via density gradient centrifugation and cryopreserved. PBMCs were thawed and rested overnight at 10^6^ cells/mL in cRPMI. The following day, samples were stimulated with 20 IU/mL rhIL-2 (Teceleukin) and/or 100 ng/mL rhIGF1 (BioVision) for 15 or 60 minutes at 37°C and then immediately fixed with an equal volume of Cytofix fixation buffer (BD Biosciences) for 10 minutes at 37°C. Live/Dead fixable near-IR dead cell stain kit (Invitrogen) was applied according to manufacturer’s instructions for dead cell exclusion. Cells were washed once with stain buffer, permeabilized with Phosflow perm buffer III (BD Biosciences) for 30 minutes at 4°C:, washed twice with stain buffer, and then incubated with human TruStain FcX (BioLegend) for 5 minutes at 23°C. Samples were stained with the following fluorescently-labeled anti-human antibodies at 23°C for 45 minutes: CD3-PerCP-Cy5.5 (UCHT1), CD45RA-BV711 (HI100), FOXP3-AF488 (206D), FOXP3-AF488 (259D), Helios-Pacific Blue (22F6) (BioLegend), CD4-PE-Cy7 (RPA-T4, eBioscience), pS6 Ser235/236-AF647 (D57.2.2E, Cell Signaling Technology), and pSTAT5 Tyr694-PE (47/Stat5, BD Biosciences). Samples were washed once with stain buffer prior to data acquisition on a Cytek Aurora 5L (16UV-16V-14B-10YG-8R) spectral flow cytometer and analysis with FlowJo software (v10.6.1; Tree Star).

### *In vitro* Homeostatic Proliferation

Naïve CD4^+^ T cells were isolated by either density gradient-centrifugation of whole blood followed by EasySep human naïve CD4^+^ T cell enrichment kit (19155, StemCell Technologies) or by RosetteSep human CD4^+^ T cell enrichment (StemCell Technologies) of whole blood followed by human CD45RO microbead depletion (Miltenyi Biotec). Enriched cells were stained with Cell Proliferation Dye eFluor670 (Thermo Fisher) according to manufacturer’s instructions. Naïve CD4^+^ T cells were plated at 10^6^ cells/mL in cRPMI with 20 IU/mL rhIL-2 (Teceleukin) and 100 ng/mL rhIGF1 (BioVision). Cytokine and/or growth factor were replenished on day 3 and day 7, assuming consumption, and intracellular flow cytometry was performed on day 9-11. Live/Dead fixable near-IR dead cell stain kit (Invitrogen) was applied according to manufacturer’s instructions for dead cell exclusion. Cells were washed once with stain buffer and incubated with human TruStain FcX (BioLegend) for 5 minutes at 4°C. Samples were stained with the following fluorescently-labeled anti-human antibodies for 30 minutes at 4°C: CD4-PerCP-Cy5.5 (RPA-T4), CD45RA-BV711 (HI100), and CD221-PE (1H7) (BioLegend) prior to fixation and permeabilization with FOXP3 transcription factor staining buffer set according to manufacturer’s protocol (eBioscience). Intracellular staining was performed with the following fluorescently-labeled anti-human antibodies for 30 minutes at 23°C: FOXP3-AF488 (206D), FOXP3-AF488 (259D), and Helios-Pacific Blue (22F6) (BioLegend). Data were acquired on an LSRFortessa (BD Biosciences) and analyzed with FlowJo software (v10.6.1; Tree Star).

### Human IL-2R Subunit and IGF1R Cross-Regulation by IGF1 and IL-2

Whole blood was processed to PBMCs via density gradient centrifugation and frozen. PBMCs were thawed and rested overnight at 10^6^ cells/mL in cRPMI. The following day, samples were stimulated with 20 IU/mL rhIL-2 (Teceleukin) and/or 100 ng/mL rhIGF1 (BioVision) and incubated for two days at 37°C. Live/Dead fixable near-IR dead cell stain kit (Invitrogen) was applied according to manufacturer’s instructions for dead cell exclusion. Cells were washed once with stain buffer and then, incubated with human TruStain FcX (BioLegend) for 5 minutes at 4°C. Samples were stained with the following fluorescently-labeled anti-human antibodies at 4°C for 30 minutes: CD4-BV711 (RPA-T4), CD45RA-BV605 (HI100), CD197-BV421 (G043H7), CD221-PE (1H7), CD25-PE-Cy5 (BC96), CD122-PerCP-Cy5.5 (TU27), CD132-APC (TUGh4) (BioLegend). Samples were fixed and permeabilized with Foxp3 transcription factor staining buffer set according to manufacturer’s protocol (eBioscience) before staining for 30 minutes at 23°C with anti-human FOXP3-AF488 (206D), FOXP3-AF488 (259D), and Helios-Pacific Blue (22F6) (BioLegend). Samples were washed once with stain buffer prior to data acquisition on a Cytek Aurora 5L (16UV-16V-14B-10YG-8R) spectral flow cytometer and analysis with FlowJo software (v10.6.1; Tree Star).

### Human T cell Transduction

Naïve CD4^+^ T cells were isolated by density gradient-centrifugation of PBMCs from whole blood followed by use of Naïve CD4^+^ T cell Isolation Kit II (Miltenyi Biotec). Cells were stained with Cell Proliferation Dye eFluor670 (eBioscience) and cultured at 10^6^/mL with 20 IU/mL rhIL-2 (Teceleukin), 10 ng/mL rhIL-7 (BD Biosciences), and/or 100 ng/mL rhIGF1 (BioVision) for 7 days prior to transduction. Cells were transduced using 8 µg/mL protamine sulfate (Sigma-Aldrich) and 3 TU/cell lentiviral vector containing R164, a T cell receptor (TCR) recognizing T1D-relevant epitope glutamic acid decarboxylase 65 (GAD65) 555-567 in the context of HLA-DRB1*04:01 (48). Lentiviral vector composition (46) and production (47) were as previously described. Cells were spinoculated by centrifugation at 1000×*g* for 30 min at 32°C (47) and IL-2, IL-7, and/or IGF1 added assuming consumption on days 3, 7, and 10. Cells were harvested on day 14, and Live/Dead fixable near-IR dead cell stain kit (Invitrogen) was applied according to manufacturer’s instructions for dead cell exclusion. Cells were washed once with stain buffer, then incubated with human TruStain FcX (BioLegend) for 5 minutes at 4°C. Samples were stained with the following fluorescently-labeled anti-human antibodies at 4°C for 30 minutes: TCR Vβ5.1-PE (IMMU 157, Beckman Coulter), CD45RA-BV711 (HI100), and CD197-PE-Cy7 (G043H7) (BioLegend). Samples were fixed and permeabilized with Foxp3 transcription factor staining buffer set according to manufacturer’s protocol (eBioscience) before staining for 30 minutes at 23°C with anti-human FOXP3-AF488 (206D), FOXP3-AF488 (259D), and Helios-Pacific Blue (22F6) (BioLegend). Samples were washed once with stain buffer prior to data acquisition on an LSRFortessa (BD Biosciences) and analysis with FlowJo software (v10.6.1; Tree Star).

### Statistics

Analyses were performed using GraphPad Prism software version 7.0. Data are presented as mean ± standard deviation (SD), and all tests were two-sided unless otherwise specified. Murine IGF1R expression was compared between cell subsets using repeated measures one-way ANOVA with Bonferroni’s multiple comparisons test. Murine and human IGF1R signaling was compared between treatment groups and timepoints using repeated measures two-way ANOVA with Tukey’s multiple comparisons test. One-way ANOVA with Dunnett’s or Tukey’s multiple comparisons tests were used to assess outcomes of *in vivo* IL-2 + IGF1 treatment when comparing all treatment groups to each other or only to PBS control, respectively. Friedman test or Kruskal-Wallis test with Dunn’s multiple comparisons test were used to compare IGF1R expression on CD4^+^ T cell subsets within human subjects or between T1D subjects and relatives, respectively. Associations between IGF1R levels and subject age were assessed via Spearman correlation. IL-2R subunit and IGF1R expression were compared between treatment conditions using Friedman test with Dunn’s multiple comparisons test. Human *in vitro* proliferation and lentiviral transduction were compared between experimental conditions via Wilcoxon test or paired t-test, depending upon if the data were normally distributed, as assessed by Shapiro-Wilk test. P-values < 0.05 were considered significant.

### Study approval

All procedures with human samples were approved by the IRB at each institution and conducted in accordance with the Declaration of Helsinki. Written informed consent was obtained from participants (or their legal guardian in the case of minors) prior to enrollment. NOD/ShiLtJ and NOD.Foxp3-GFP/cre mice were bred at UF for *in vitro* experiments or purchased from Jax by the University of Miami (UM) for *in vivo* experiments, and housed in specific pathogen-free facilities, with food and water available *ad libitum*. All murine studies were conducted in accordance with protocols approved by the UF or UM Institutional Animal Care and Use Committee (IACUC) and in accordance with the National Institutes of Health Guide for Care and Use of Animals.

## Author Contributions

MRS designed the studies, conducted experiments, acquired data, analyzed data, and wrote the manuscript.

LDP designed the studies, conducted experiments, acquired data, analyzed data, and reviewed/edited the manuscript.

MEB designed the studies, conducted experiments, acquired data, analyzed data, and reviewed/edited the manuscript.

CCK conducted experiments, acquired data, and reviewed/edited the manuscript. ALP contributed to discussion and reviewed/edited the manuscript.

ALB designed the studies, analyzed data, and reviewed/edited the manuscript.

TMB conceptualized the project, designed studies, provided reagents, acquired funding, and reviewed/edited the manuscript.

## GRAPHICAL ABSTRACT

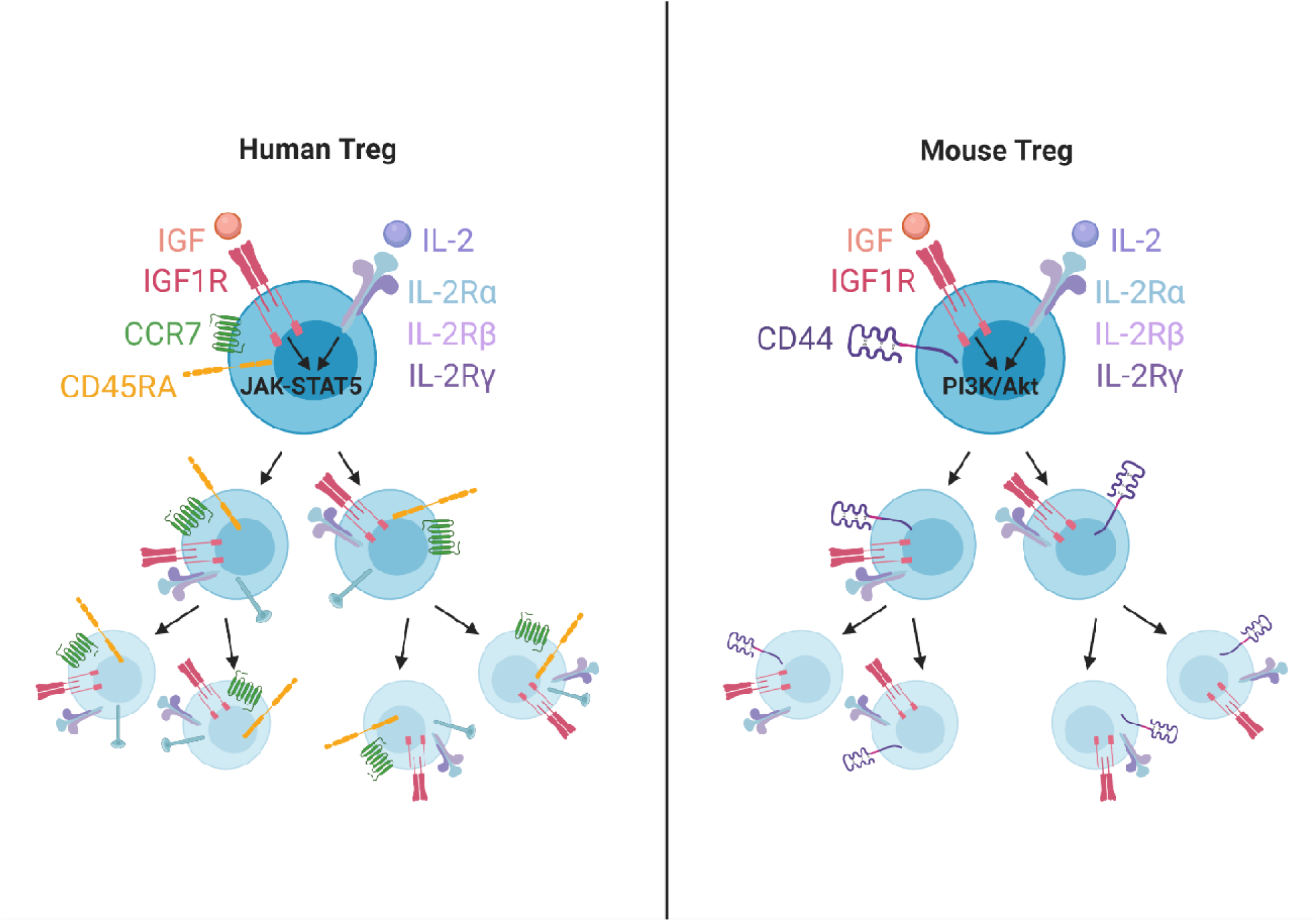

