## Supplementary Figures for "Insulin-like Growth Factor-1 Synergizes with IL-2 to Induce Homeostatic Proliferation of Regulatory T cells"

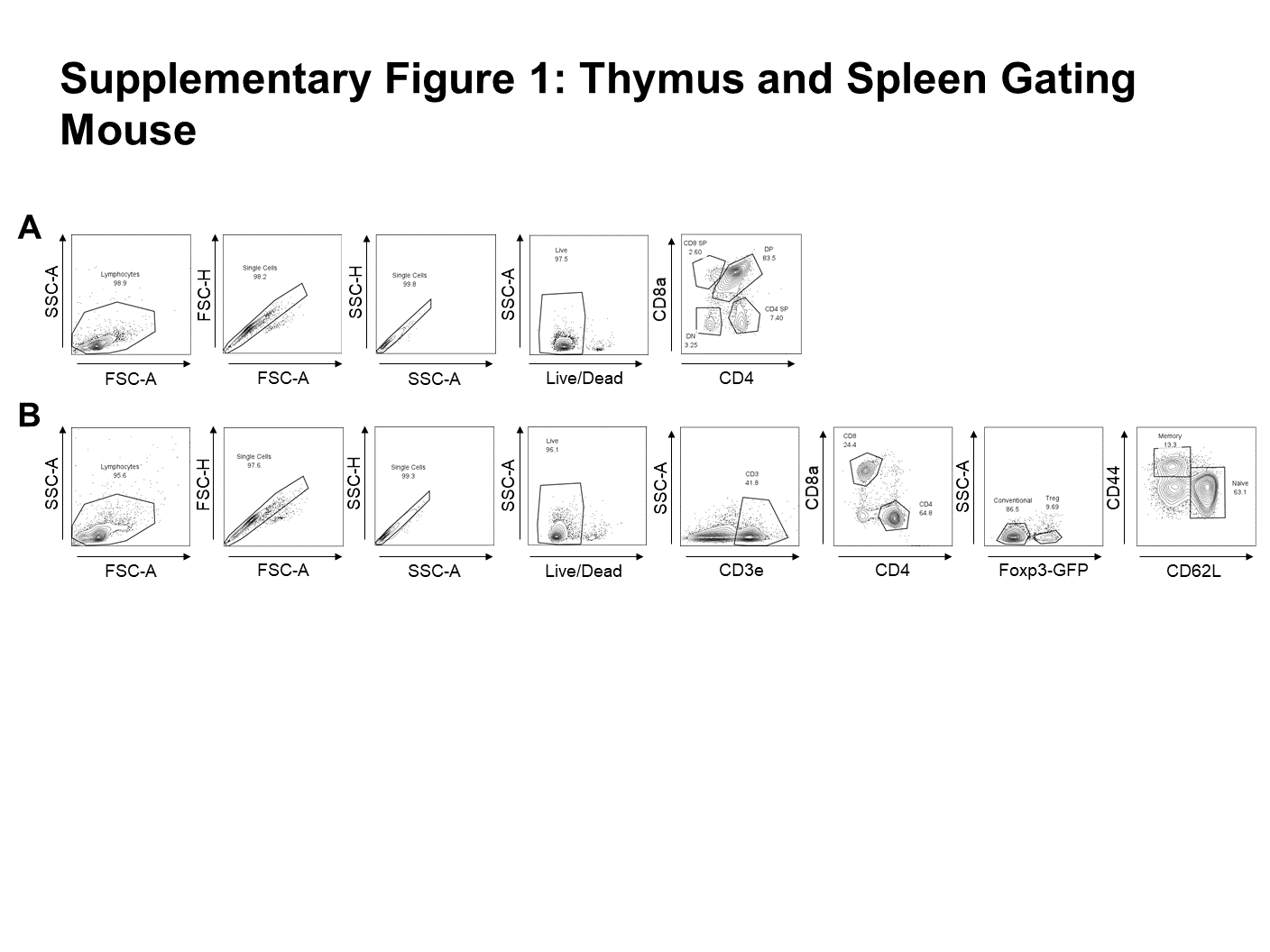

**Supplementary Figure 1. Gating strategies for murine IGF1R expression on thymocytes and CD4^+^ T cells.** Representative gating for (**A**) thymus and (**B**) spleen.

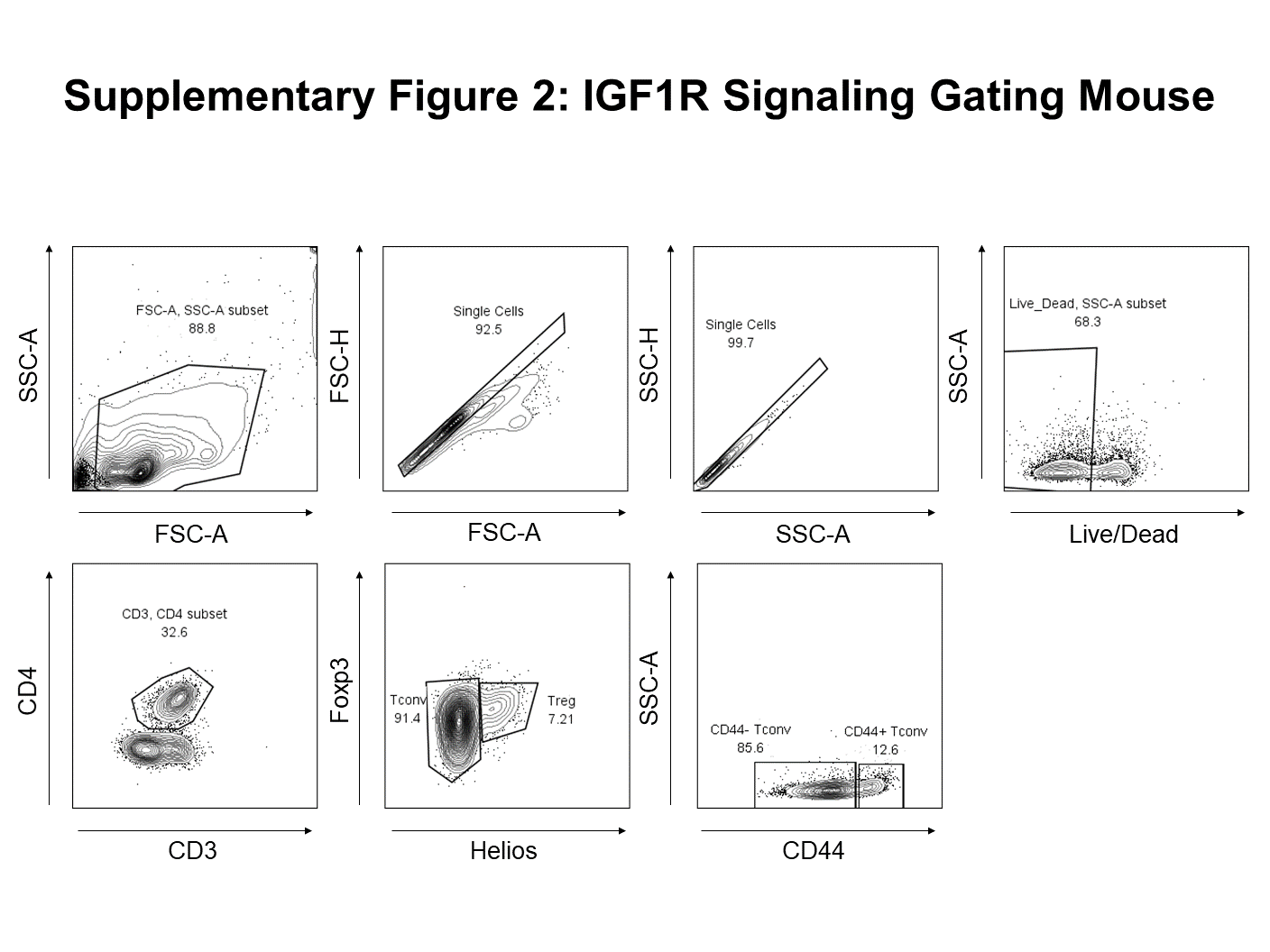

**Supplementary Figure 2. Gating strategy for murine IGF1R and IL-2R signaling.**

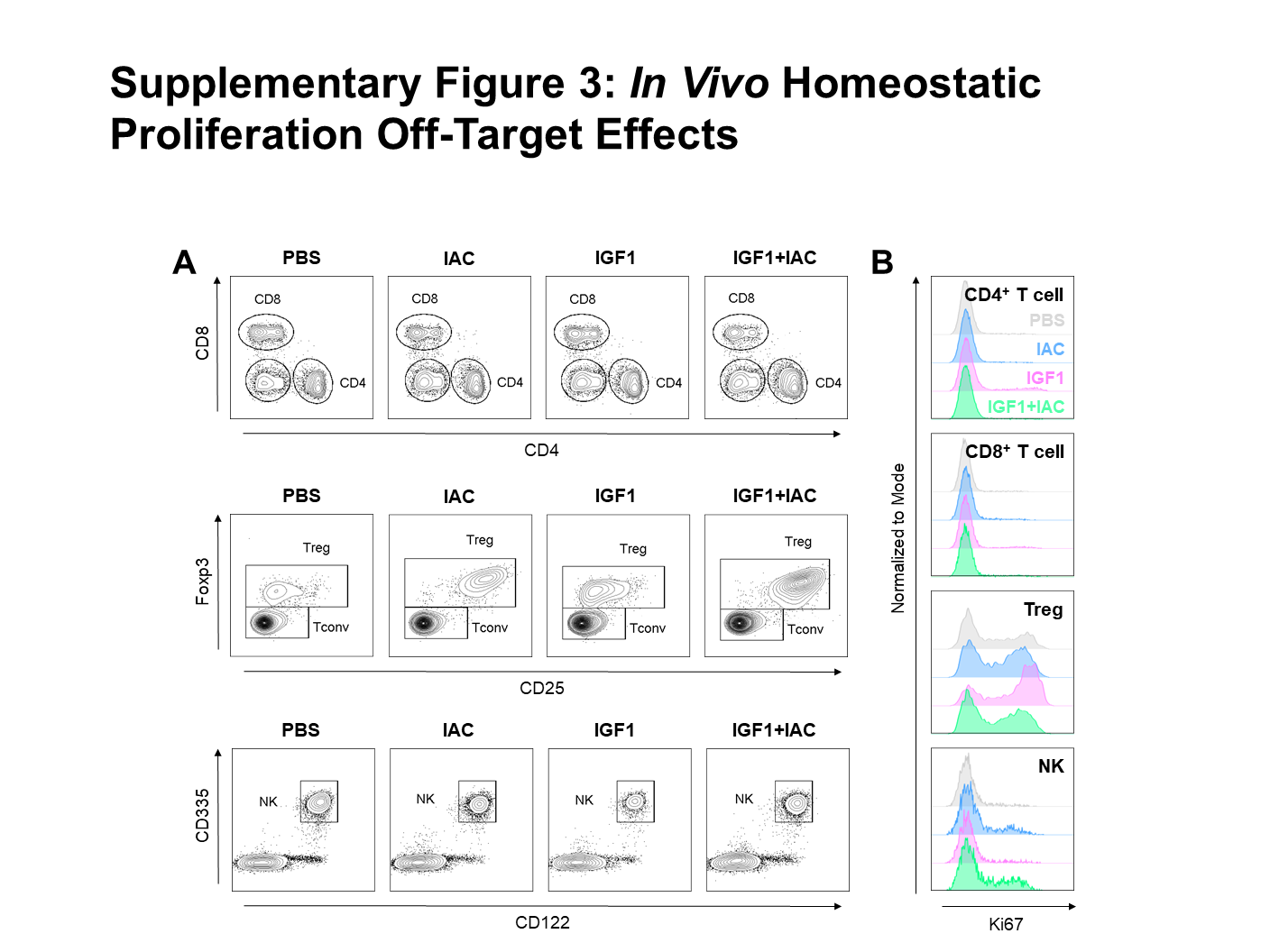

**Supplementary Figure 3. Off-target effects of IGF1 + IL-2 antibody complex (IAC) treatment in NOD mice.** (**A**) Representative contour plots showing gating of CD4^+^, CD8^+^ T cells, and CD4^+^CD25^+^Foxp3^+^ Tregs (pre-gated on CD19^-^Ly6G^-^ cells), as well as CD335^+^CD122^+^ NK cells (pre-gated on CD19^-^Ly6G^-^CD4^-^CD8^-^ cells). (**B**) Representative histograms of Ki67 expression in each cell subset in mice treated for one week with PBS (gray), IAC (blue), IGF1 (purple), or IGF1 + IAC (green).

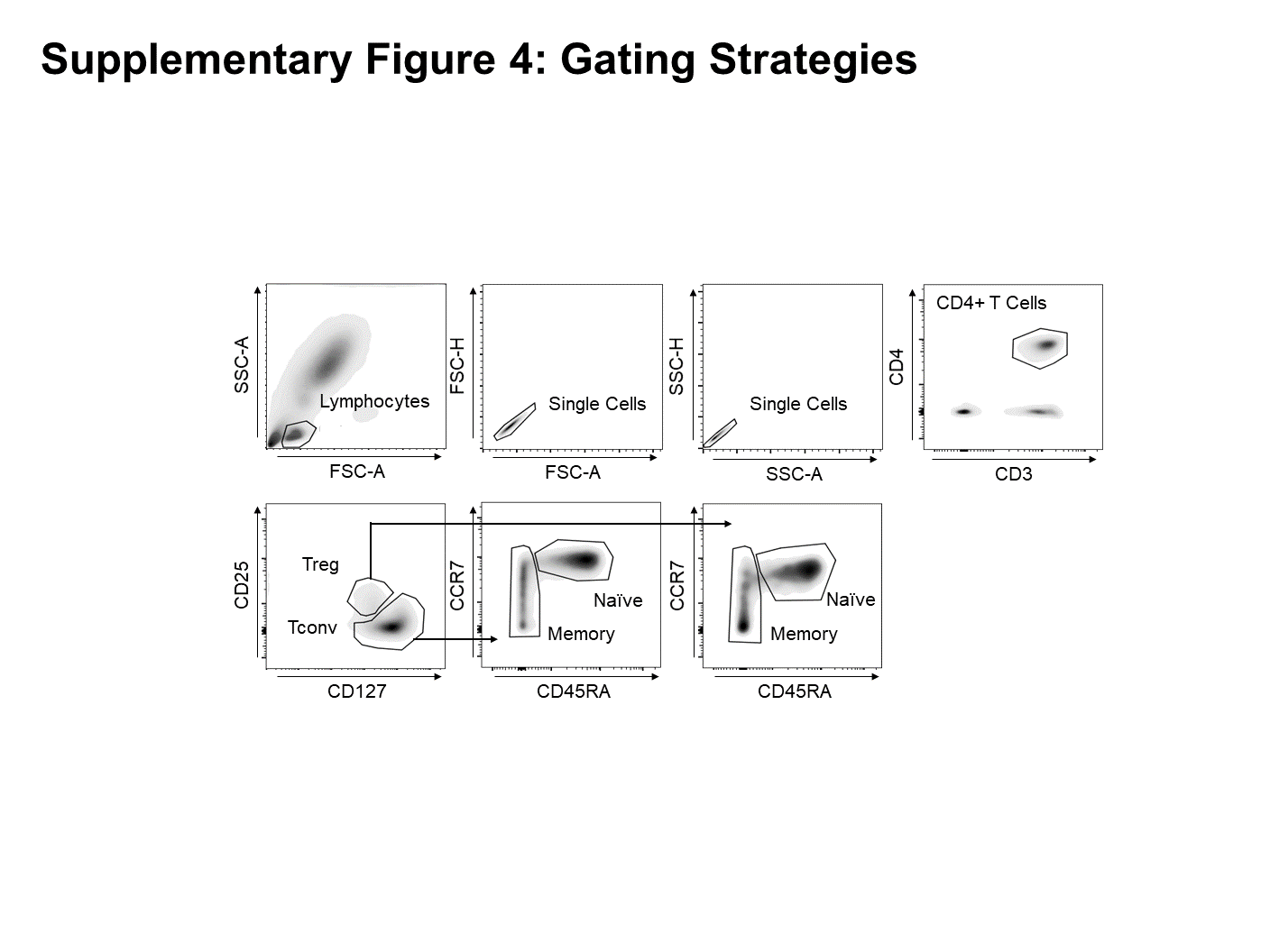

**Supplementary Figure 4. Gating strategy for human IGF1R expression on CD4^+^ T cells.**

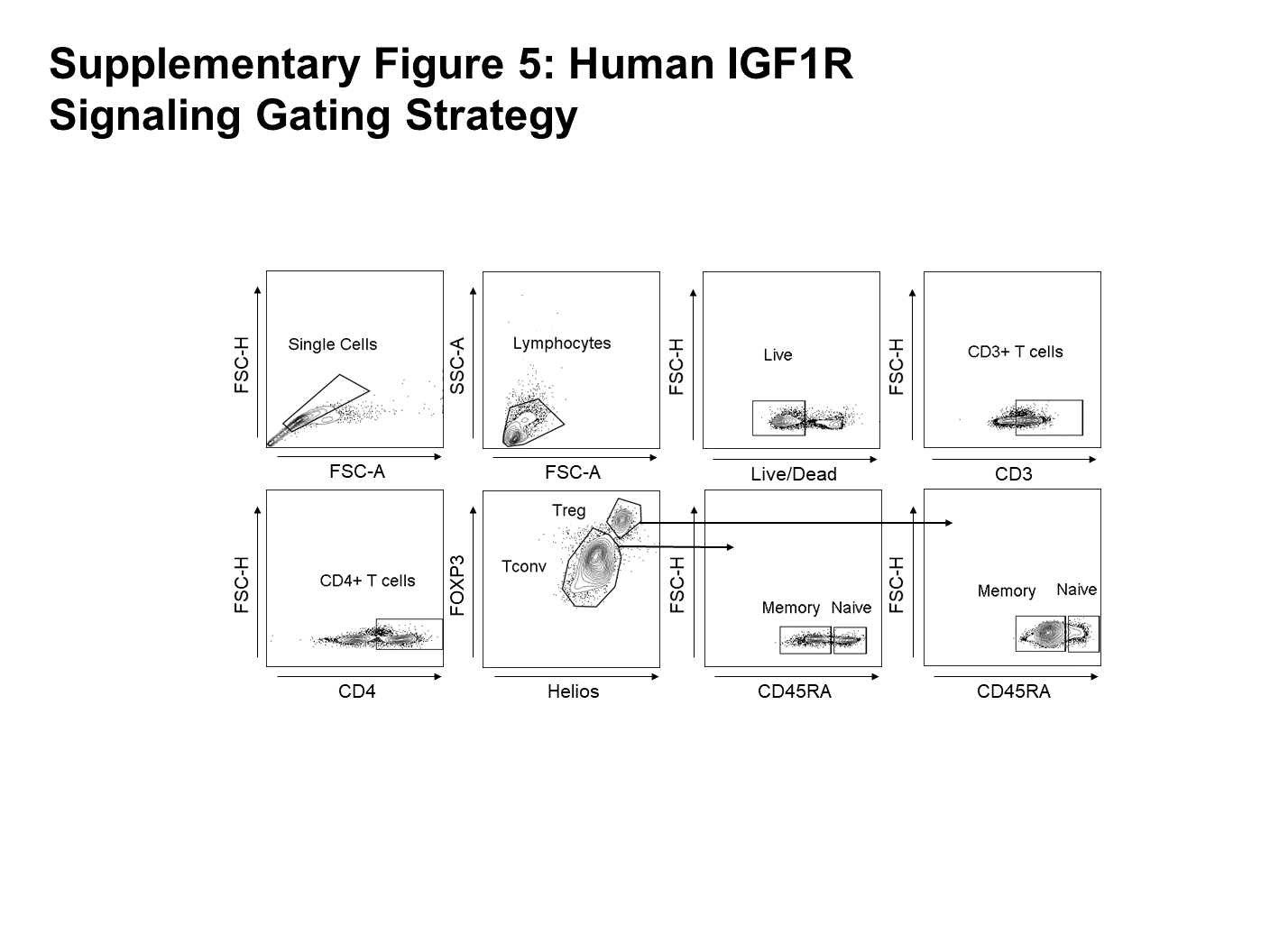

**Supplementary Figure 5. Gating strategy for human IGF1R and IL-2R signaling.**

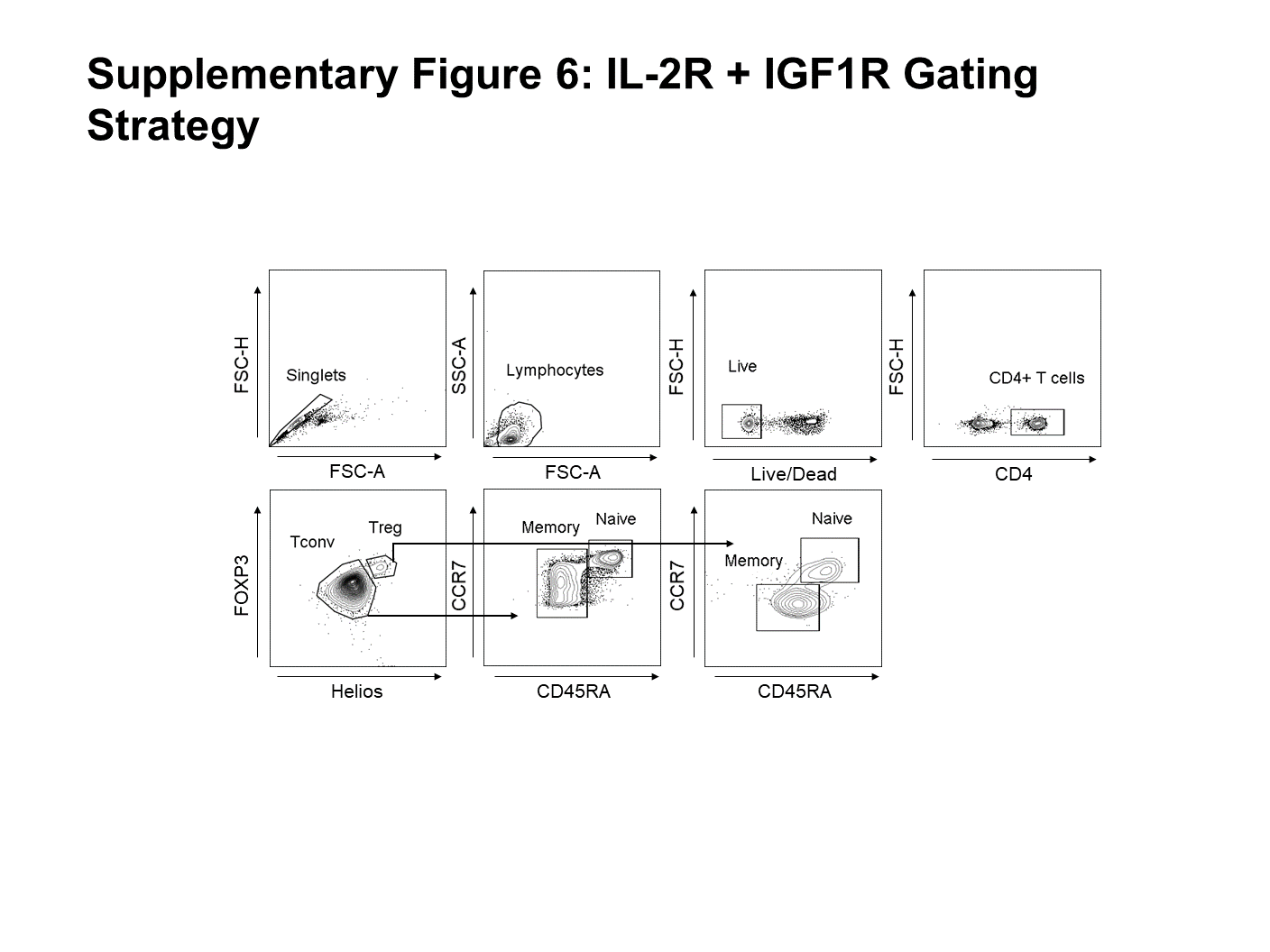

**Supplementary Figure 6. Gating strategy for human IGF1R and IL-2R subunit expression**

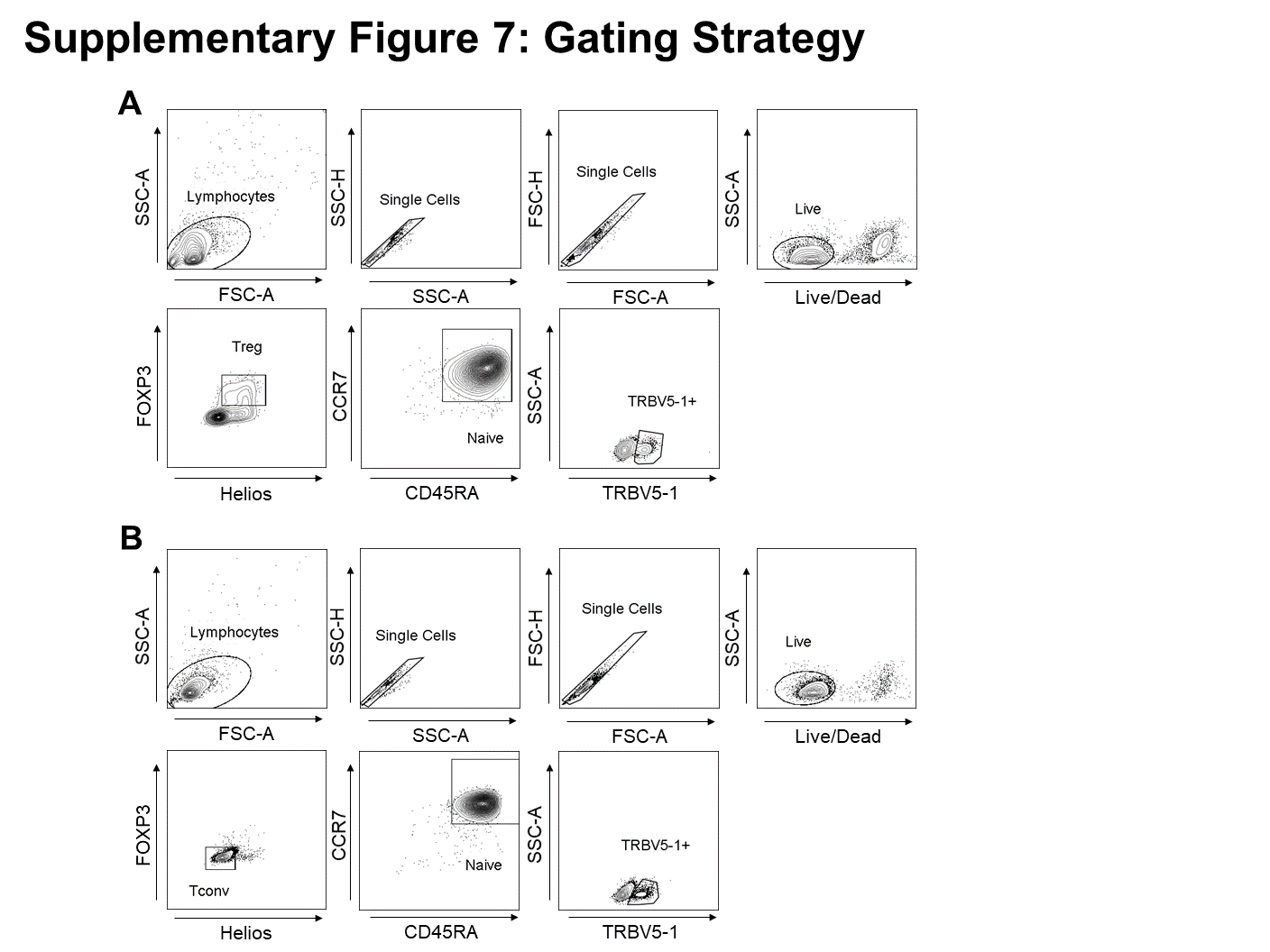

**Supplementary Figure 7. Gating strategy for homeostatic transduction of naïve human CD4^+^ T cells.** Representative gating for transduction of (**A**) naïve Tregs using IGF1 + IL-2 or (**B**) naïve Tconv using IGF1 + IL-7.
